# A Neuropsin-based Optogenetic Tool for Precise Control of G_q_ signaling

**DOI:** 10.1101/2022.02.22.481462

**Authors:** Ruicheng Dai, Tao Yu, Danwei Weng, Heng Li, Yuting Cui, Zhaofa Wu, Qingchun Guo, Haiyue Zou, Wenting Wu, Xinwei Gao, Zhongyang Qi, Yuqi Ren, Shu Wang, Yulong Li, Minmin Luo

**Affiliations:** School of Life Sciences, Tsinghua University, Beijing 100084, China; National Institute of Biological Sciences (NIBS), Beijing 102206, China; School of Life Sciences, Peking University, Beijing 100871, China; Graduate School of Peking Union Medical College, Beijing 100730, China; Chinese Institute for Brain Research, Beijing 102206, China; Tsinghua Institute of Multidisciplinary Biomedical Research (TIMBR), Beijing 102206, China; Peking University-Tsinghua University-NIBS Joint Graduate Program, NIBS, Beijing 102206, China; State Key Laboratory of Membrane Biology, Peking University School of Life Sciences, Beijing 100871, China; PKU-McGovern Institute for Brain Research, Beijing 100871, China; Capital Medical University, Beijing 102206, China; Academy for Advanced Interdisciplinary Studies, Peking University, Beijing 100871, China

## Abstract

G_q_-coupled receptors regulate numerous physiological processes by activating enzymes and inducing intracellular Ca^2+^ signals. There is a strong need for an optogenetic tool that enables powerful experimental control over G_q_ signaling. Here, we present chicken opsin 5 (cOpn5) as the long sought-after, single-component optogenetic tool that mediates ultra-sensitive optical control of intracellular G_q_ signaling with high temporal and spatial resolution. Expressing cOpn5 in mammalian cells enables blue light-triggered, G_q_-dependent Ca^2+^ release from intracellular stores and protein kinase C activation. Strong Ca^2+^ transients were evoked by brief light pulses of merely 10 ms duration and at 3 orders lower light intensity of that for common optogenetic tools. Photostimulation of cOpn5-expressing cells at the subcellular and single-cell levels generated intracellular and intercellular Ca^2+^ wave propagation, respectively, thus demonstrating the high spatial precision of cOpn5 optogenetics. The cOpn5-mediated optogenetics could also be applied to activate neurons and control animal behavior in a circuit-dependent manner. We further revealed that optogenetic activation of cOpn5-expressing astrocytes induced massive ATP release and modulation neuronal activation in the brain of awake, behaving mice. cOpn5 optogenetics may find broad applications in studying the mechanisms and functional relevance of G_q_ signaling in both non-excitable cells and excitable cells in all major organ systems.

## Main

G-protein-coupled receptors (GPCRs) modulate many intracellular signaling pathways and represent some of the most intensively studied drug targets^1^. Upon ligand binding, the GPCR undergoes a conformation change that is transmitted to heterotrimeric G proteins, which are multi-subunit complexes comprising G_α_ (G_q/11_, G_s_, G_i/o_ and G_12/13_) and tightly associated G_βγ_ subunits^2^. The G_q_ proteins, a subfamily of G_α_ subunits, couple to a class of GPCRs to mediate cellular responses to neurotransmitters, sensory stimuli, and hormones throughout the body^3, 4^. Their primary downstream signaling targets include phospholipase C beta (PLC-β) enzymes, which catalyze the hydrolysis of phospholipid phosphatidylinositol bisphosphate (PIP_2_) into inositol trisphosphate (IP_3_) and diacylglycerol (DAG). IP_3_ triggers intracellular Ca^2+^ release, which together with DAG activates protein kinase C (PKC)^5^. Studies have demonstrated that the activation of G_q_-coupled receptors often produces rapid (within seconds) and robust increase in Ca^2+^ signals^6^. Given that Ca^2+^ signals and PKC activity impact nearly every cellular process in both non-excitable cells and excitable cells, developing tools that precisely control intracellular G_q_ signaling would be highly valuable for studying the mechanisms and physiological functions of G_q_-coupled receptors.

Optogenetics uses light-responsive proteins to achieve optically-controlled perturbation of cellular activities with genetic specificity and high spatiotemporal precision^7, 8^. Since the early discoveries of optogenetic tools using light-sensitive ion channels and transporters, diverse technologies have been developed and now support optical interventions into intracellular second messengers, protein interactions and degradation, and gene transcription^9–12^. Optogenetic tools that control G_q_ signaling may have several advantages over current approaches including chemogenetics and photoactivatable small molecules^13–15^: unlike chemogenetics, they potentially offer subsecond temporal resolution and subcellular spatial resolution; unlike photoactivatable compounds, they provide control of genetically-identified cell types in complex organ systems. However, despite several intensive efforts, to date there has been few optogenetic tool that enables rapid and effective activation of G_q_ signaling^16–18^. Opto-a1AR, a creatively designed G_q_-coupled rhodopsin-GPCR chimera, induces mild intracellular Ca^2+^ increase only after long-time (>60 s) photostimulation^19^. Melanopsin (Opn4) in a subset of mammalian retinal ganglion cells is a G_q_-coupled opsin that mediates no-image-forming visual functions^20–24^. However, cells expressing mOpn4L show weak Ca^2+^ responses even after prolonged (15-60 s) exposure to bright light illumination, and require continuous chromophore addition in the culture medium^25–27^. Indeed, systematic characterizations have revealed major limitations of these two tools for *in vitro* and *in vivo* applications^28–30^. Opto-a1AR and Opn4 thus only partially mimick the activation of endogenous G_q_-coupled receptors.

We asked whether some naturally occurring photoreceptors could serve as efficient optogenetic tools for G_q_ signaling. Most animals detect light using GPCR-based photoreceptors, which comprise a protein moiety (opsin) and a vitamin A derivative (retinal) that functions as both a ligand and a chromophore^31^. Several thousand opsins have been identified to date^32, 33^, and two recent studies reported G_i_-based opsins from mosquito and lamprey for presynaptic inhibition in neurons^34, 35^. Opn5 (neuropsin) and its orthologs in many vertebrates have been reported as an ultraviolet (UV)-sensitive opsin that couples to G_i_ proteins^36–39^. Interestingly, exposure to blue light induces an increase in intracellular Ca^2+^ levels within avian primary Müller glial cells endogenously expressing Opn5^40^, hinting the possibility that certain Opn5 might also couple to G_q_ proteins^41, 42^.

Here we report that the chicken Opn5 (cOpn5 for simplicity), but not two of its mammalian orthologs, sensitively and strongly mediated blue light-induced activation of G_q_ signaling in mammalian cells. Detailed characterizations of cOpn5 reveal that it is at least 3 orders of magnitude higher in light sensitivity and temporal precision than existing G_q_-coupled opsin-based tools — opto-a1AR and Opn4, provides subcellular spatial resolution, and does not require chromophore addition. We further demonstrate cOpn5 optogenetics as a highly effective approach for activating neurons to produce robust behavior changes in freely moving mice, as well as for activating astrocytes to induce massive ATP release *in vivo*. These findings establish that cOpn5 can be utilized as powerful, single-component optogenetic tools to support experimental investigations into the mechanisms and functions associated with G_q_ signaling in both non-excitable cells and excitable cells.

## Results

### cOpn5 mediates optogenetic activation of G_q_ signaling

We tested whether heterologous expression of the Opn5 orthologs from chicken, humans, and mice (which share 80-90% protein sequence identity) have the capacity to mediate blue light-induced G_q_ signaling activation within HEK 293T cells (Fig. 1a and Supplementary Table 1). The chicken Opn5 (cOpn5) was co-localized with the EGFP-CAAX membrane maker, indicating that it was effectively expressed on the plasma membrane (Fig. 1b). We used blue light for stimulation and the red intracellular Ca^2+^ indicator Calbryte^TM^ 630 AM dye to monitor the relative Ca^2+^ response (Fig. 1c; See Supplementary Table 2 for the list of resources). cOpn5 mediated an immediate and strong light-induced increase in Ca^2+^ signal (∼3-fold increase in Ca^2+^ indicator fluorescence intensity relative to its resting fluorescence intensity; abbreviated as *ΔF/F*), whereas no light effect was observed from cells expressing the human or mouse Opn5 orthologs (Fig. 1d; Video 1). Note that we did not supply any exogenous retinal to the culture media, which suggested that endogenous retinal is sufficient to render cOpn5 functional^43^. Thus, unlike mammalian Opn5, cOpn5 allows effective optical activation of intracellular Ca^2+^ signals in the human HEK 293T cells.

**Fig. 1:**
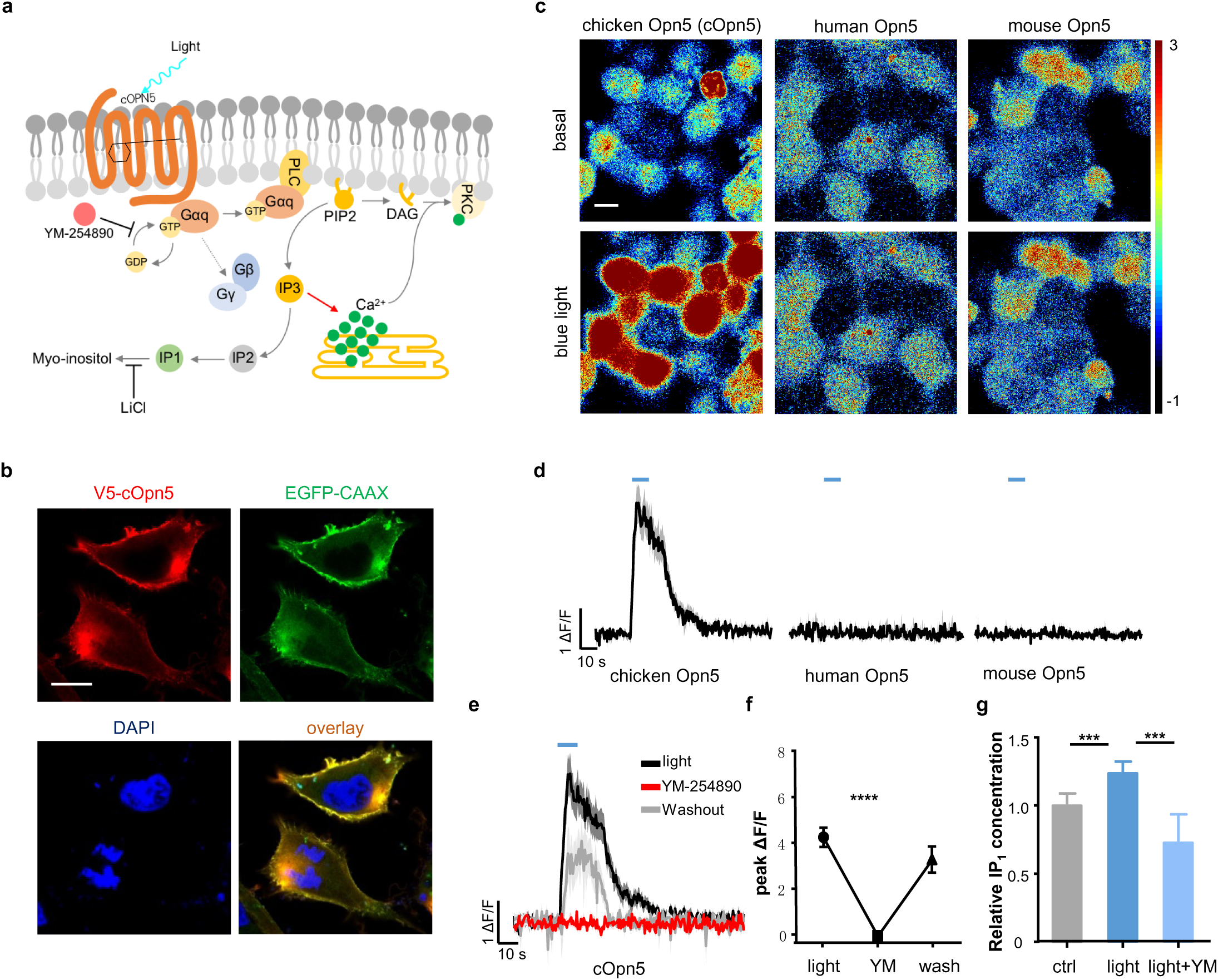
cOpn5 mediates optical activation of G_q_ signaling in HEK 293T cells. **a**, Schematic diagram of the putative intracellular signaling in response to light-induced cOpn5 activation. PLC: phospholipase C; PIP_2_: phosphatidylinositol-4,5-bisphosphate; IP_3_: inositol-1,4,5-trisphosphate; IP_1_: inositol monophosphate; DAG: diacylglycerol; PKC: protein kinase C; YM-254890: a selective G_q_ protein inhibitor. **b**, The Cy3-counterstained V5-cOpn5 fusion protein (red) was co-localized with the membrane-tagged EGFP-CAAX (green) in HEK 293T cells. DAPI counterstaining (blue) indicates cell nuclei. Scale bar, 10 μm. **c**, Pseudocolor images of the Ca^2+^ signals before and after blue light stimulation (10 s; 100 μW/mm^2^; 488 nm) in HEK 293T cells expressing Opn5 from three species (*Gallus gallus, Homo sapiens, and Mus musculus*). Scale bar, 10 μm. **d**, Time courses of light-evoked Ca^2+^ signals for cells shown in **c**. Blue lines above the curves indicate light stimulation. Ca^2+^ indicator fluorescence intensity relative to its resting fluorescence intensity; abbreviated as ΔF/F. **e**, The G_q_ protein inhibitor YM-254890 (10 nM) reversibly blocked cOpn5-mediated, light-induced Ca^2+^ signals (n = 28 HEK 293T cells). **f**, Group data show that the G_q_ protein inhibitor YM-254890 (10 nM) reversibly blocked cOpn5-mediated, light-induced Ca^2+^ signals (N = 28 HEK 293T cells). ****P <0.0001, one way ANOVA. Error bars indicate S.E.M. **g**, YM suppressed the IP_1_ accumulation evoked by continuous light stimulation (3 min; 100 μW/mm^2^; 470 nm) in cOpn5-expressing HEK 293T cells (N = 4 replications). ***P < 0.005, unpaired t tests).

We then examined whether cOpn5 truly mediated the activation of G_q_ signaling, which triggers signal cascades that produce two second messengers: IP_3_ leading to Ca^2+^ releases from intracellular stores, and DAG leading to PKC activation. Preincubation of YM-254890, a highly selective G_q_ proteins inhibitor^44^, reversibly abolished the light-induced Ca^2+^ transients in both cOpn5-expressing cells (Figs. 1e and 1f). We observed strong Ca^2+^ signals in the absence of extracellular Ca^2+^, thus indicating Ca^2+^ release from intracellular stores (Extended Data Fig. 1a). In cOpn5-, but not human OPN5-expressing cells, we also detected a light-induced increase in the level of inositol phosphate (IP_1_), the rapid degradation product of IP_3_; moreover, the extent of this increase was reduced with the treatment of YM-254890 (Fig.1g and Extended Data Fig. 1d; See Supplementary Table 3 for detailed information of statistical analyses). This result thus indicated light-evoked, G_q_-dependent IP_3_ production. In cOpn5-expressing HEK 293T cells, blue light also triggered the phosphorylation of MARCKS protein, a well-established target of PKC^45^, in a PKC activity-dependent manner (Extended Data Figs. 1b and 1c). Consistent with earlier findings indicating that mammalian Opn5 couples to G_i_^36–38^, blue light illumination effectively reduced cAMP levels in cells expressing human and mouse Opn5 with retinal addition; however, it had very mild effect in cOpn5-expressing cells in the presence of retinal and had no effect without retinal (Extended Data Fig. 1e). Collectively, these data revealed that blue light illumination enables the efficient coupling of cOpn5, but not its mammalian orthologs, to the G_q_ signaling pathway in mammalian cells.

### cOpn5-mediated optogenetics is sensitive and precise

Given that light-stimulated cOpn5 mimics endogenous G_q_-coupled GPCRs to rapidly activate the G_q_ signaling pathway, we asked whether cOpn5 could serve as the long sought-after optogenetic tool for G_q_ signaling, and more importantly, whether it has features common to other popular optogenetic tools, such as high light sensitivity, single-component convenience, and high spatiotemporal resolution. We first systematically characterized the light-activating properties of cOpn5-expressing HEK 293T cells. Although Opn5 is previously considered as an ultraviolet (UV)-sensitive photoreceptor^37^, mapping with a set of wavelengths ranging 365-630 nm at a fixed light intensity of (100 μW/mm^2^) revealed that the 470 nm blue light elicited the strongest Ca^2+^ transients, with the UVA light (365 and 395 nm) being less effective and longer-wavelength visible light (561 nm or above) completely ineffective (Fig. 2a). We then tested the effects of varying photostimulation durations. Stimulating with brief light pulses (1, 5, 10, 20, 50 ms; 16 μW/mm^2^; 470 nm) showed that the Ca^2+^ response achieved the saturation mode with light duration over 10 ms. Longer light durations did not further increase the Ca^2+^ signal amplitude at this light intensity (Fig. 2b). Delivering brief blue light pulses (10 ms; 470 nm) at different intensities showed that the light of ∼4.8 μW/mm^2^ and 16 μW/mm^2^ produced about half maximum and full maximum responses, respectively (Fig. 2c). The response time courses revealed that 10 ms blue light pulses (16 μW/mm^2^) generated significant Ca^2+^ signals within 1 s and produced peak responses within 2.5 s (Fig. 2c). For 10 ms, 16 μW/mm^2^ blue light stimulation, time to 10% peak activation was 1.36 ± 0.55 s; time to 90% peak activation was 2.37 ± 0.87 s; decay time τ = 18.66 ± 4.98 s. Therefore, the light sensitivity of cOpn5 is 3-4 orders of magnitude higher than the reported values of the light-sensitive G_q_-coupled GPCRs and even 2-3 orders higher than those of the commonly used optogenetic tool Channelrhodopsin-2 (ChR2)^46, 47^ (Supplementary Table 4). Together, these results indicate that cOpn5 could function as a single-component optogenetic tool without additional retinal, and that cOpn5 is super-sensitive to blue light for its full activation requiring low light intensity (16 μW/mm^2^) and short duration (10 ms).

**Fig. 2:**
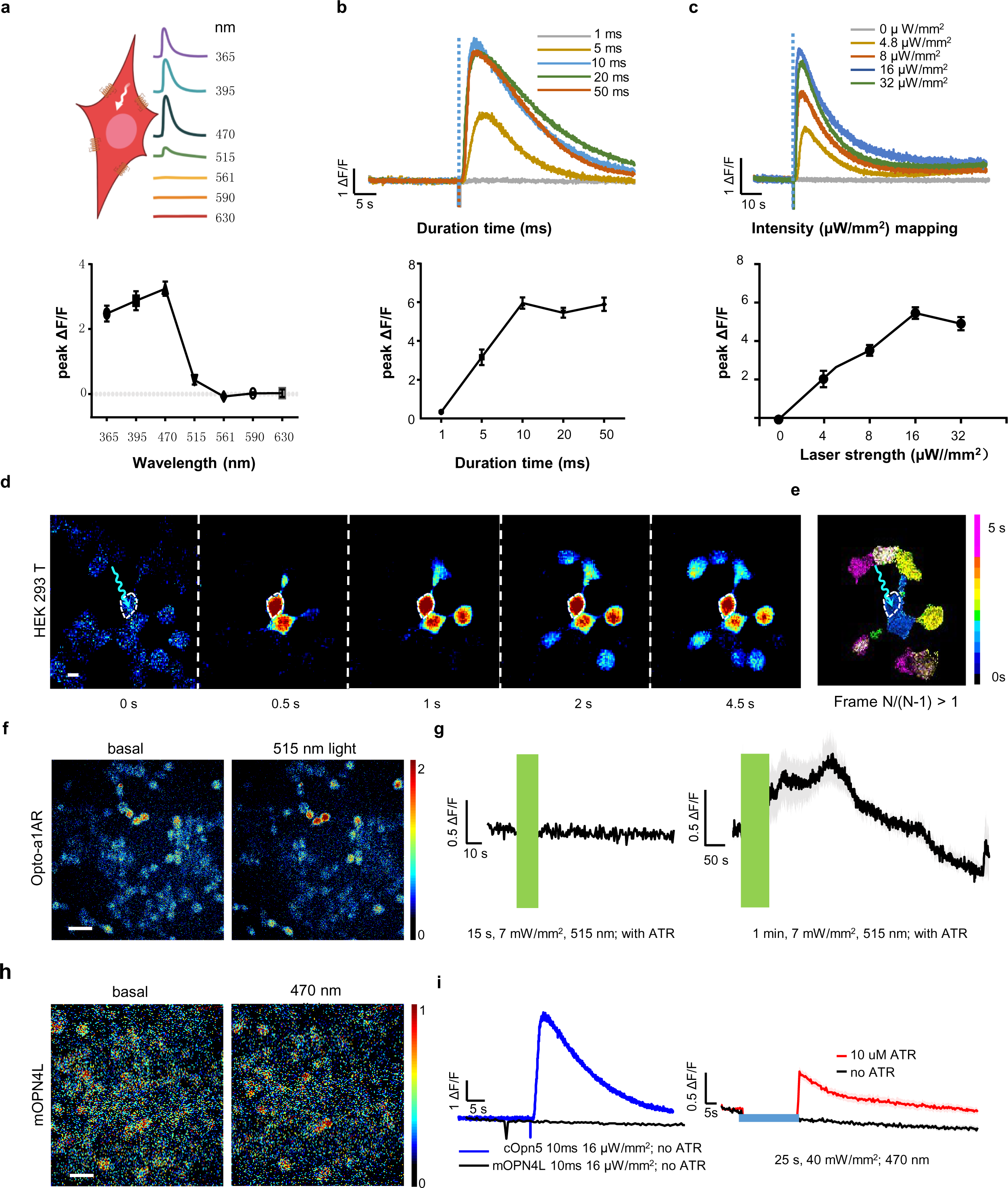
cOpn5 sensitively mediates optical control of G_q_ signaling with high temporal and spatial resolution. **a**, Schematic diagram of selected wavelengths (365, 395, 470, 515, 561, 590, and 630 nm) and the raw traces of Ca^2+^ signals (top panel) and the mean amplitudes of Ca^2+^ signal of cOpn5-expressing HEK 293T cells in response to light stimulation with different wavelengths (2 s; 100 μW/mm^2^; bottom panel). Error bars indicate S.E.M.. **b**, Time course of Ca^2+^ signals evoked by cOPN5-mediated optical activation using light pulses of different durations (1, 5, 10, 20, or 50 ms; 16 μW/mm^2^; 470 nm; N = 49 HEK 293T cells) (top panel) an the mean response magnitudes to light stimulation of different durations (1, 5, 10, 20, or 50 ms; 16 μW/mm^2^; 470 nm) (bottom panel). Error bars indicate S.E.M.. **c**, Time course of cOpn5-mediated Ca^2+^ signals under different light intensity (0, 4.8, 8, 16, or 32 μW/mm^2^; 10 ms; 470 nm, mean ± S.E.M.; N = 10 cells) (top panel) and the mean response magnitude under different light intensities (0, 4.8, 8, 16, or 32 μW/mm2) at 10 ms, 470 nm (n = 88 HEK 293T cells) (bottom panel). Error bars indicate S.E.M.. For 10 ms,16 μW/mm2 stimulation, time to 10% peak activation = 1.36 ± 0.55 s; time to 90% peak activation = 2.37 ± 0.87 s; decay time τ = 18.66 ± 4.98 s. **d**, Images of light-induced (63 ms; 17 μW; arrow points to the stimulation region) Ca^2+^ signal propagation from the stimulated HEK 293T cell to surrounding cells. Scale bar, 10 μm. **e**, Pseudocolor images showing the process of Ca^2+^ signal propagation across time of **d** (frame N/(N-1) > 1). Frame interval was 500 ms and each frame is counted once. **f**, Pseudocolor images of the baseline and peak Ca^2+^ signals (*ΔF/F0*) in opto-a1AR-expressing HEK 293T cells. The medium buffer contains 10 μM all-trans-retinal. Scale bar, 30 μm. **g**, The lack of effect by 15 s light stimulation on Ca^2+^ signals (left panel) and mild effect of 60 s light stimulation on the Ca^2+^ in opto-a1AR-expressing HEK 293T cells (right panel; N = 15 cells). Green bars indicate light stimulations. **h**, Pseudocolor images of the baseline and peak Ca^2+^ signals (*ΔF/F0*) in mOPN4L-expressing HEK 293T cells. The medium buffer contains 10 μM all-trans-retinal. Scale bar, 30 μm. **i**, The left panel shows that without ATR, brief light pulses (10 ms, 16 μW/mm^2^, 470 nm) evoked strong Ca^2+^ signals in cOpn5-expressing cells (blue line; N = 10 HEK 293T cells) but had no effect on mOPN4L-expressing cells (black line; N =12 HEK 293T cells). The right panel shows the effect of 25 s, 40 mW/mm^2^ light stimulation on the Ca^2+^ in mOPN4L-expressing HEK 293T cells within 10 μM ATR (N = 12 cells; red line) and the lack of such effect following ATR removal (black line).

Designer Receptors Exclusively Activated by Designer Drugs (DREADD)-based chemogenetic tools efficiently modulate cellular activity^48, 49^. For example, hM3Dq expression allows the activation of G_q_ signaling by adding the exogenous small molecule ligand clozapine-N-oxide (CNO)^14, 50, 51^. We thus compared the Ca^2+^ signals in response to 10 s infusion of CNO on hM3Dq-expressing HEK 293T cells to that evoked by 10 s photostimulation of cOpn5-expressing cells. Although the response amplitudes were similar, cOpn5-mediated optogenetic stimulation produced faster and temporally more precise response, as well as more rapid recovery than hM3Dq-mediated chemogenetic stimulation (Fig. 2b and Extended Data Fig. 2a-c). These results thus indicate that cOpn5-mediated optogenetics are more controllable in temporal accuracy than those of hM3Dq-mediated chemogenetics.

What’s more, cOpn5 optogenetics provides the major advantage of spatially precise control of cellular activity. Restricting brief light stimulation (63 ms) into individual cOpn5-expressing HEK 293T cells resulted in the immediate activation of the stimulated cell. Interestingly, in high cell confluence areas, the Ca^2+^ signals propagated to surrounding cells, thus suggesting intercellular communication among HEK 293T cells through a yet-to-identified mechanism (Fig. 2d, 2e and Supplementary Video 2).

We directly compared the performance of cOpn5 to those of opto-a1AR and Opn4, which had been proposed for optogenetic control of G_q_ signaling. Following the protocol in a previous report^19^, we found that 15 s illumination at the rather strong intensity level (7 mW/mm^2^) was completely ineffective in opto-a1AR-epxpressing cells; further increasing the light exposure duration to 60 s triggered a slow and small (∼0.5 *ΔF/F*) Ca^2+^ signal increase (Figs. 2f and 2g). We also compared the performance of cOpn5 to that of mouse Opn4L, a natural opsin which was reported as a tool for G_q_ signaling activation^26, 52^. Without the addition of all-trans-retinal (ATR), neither brief blue light pulses (16 μW/mm^2^) nor prolonged strong light illumination (25 s, 40 mW/mm^2^, 470 nm) had slightly effect on the Ca^2+^ signals in mOPN4L-expressing HEK 293T cells (Figs. 2h and 2i). Following the addition of exogenous ATR, long exposure of very strong illumination (25 s; 40 mW/mm^2^) triggered a slow Ca^2+^ signal increase (∼1 *ΔF/F*) in mOPN4L-expressing cells; by contrast, in cOpn5-expressing cells the light pulses of only 1/2500 duration (10 ms) and 1/2500 intensity (16 μW/mm^2^) produced nearly 6-fold increase in Ca^2+^ signals (Figs. 2h and 2i). Therefore, compared with existing opsin-based tools (opto-a1AR and mOpn4L), cOpn5 is much more light-sensitive (at least ∼3 orders higher sensitivity), requires much shorter time exposure (10 ms vs. 25 s or 60 s), and produces severalfold stronger responses (Supplementary Table 4).

### cOpn5 optogenetics activates neurons and modulates animal behaviors

We explored the application of cOpn5-mediated optogenetics directly in neurons. We first examined whether cOpn5 could mediate light-induced Ca^2+^ signals. Using AAV and the pan-neuronal SYN promoter, we expressed cOpn5 and the red Ca^2+^ indicator jRGECO1a in mouse cortical neurons (Fig. 3a). In brain slice preparations, application of blue light pulses (10s; 100 μW/mm^2^; 473 nm) reliably evoked Ca^2+^ transients in neurons (Figs. 3b and 3c). Thus, cOpn5 also enables light-induced activation in neurons.

**Fig. 3:**
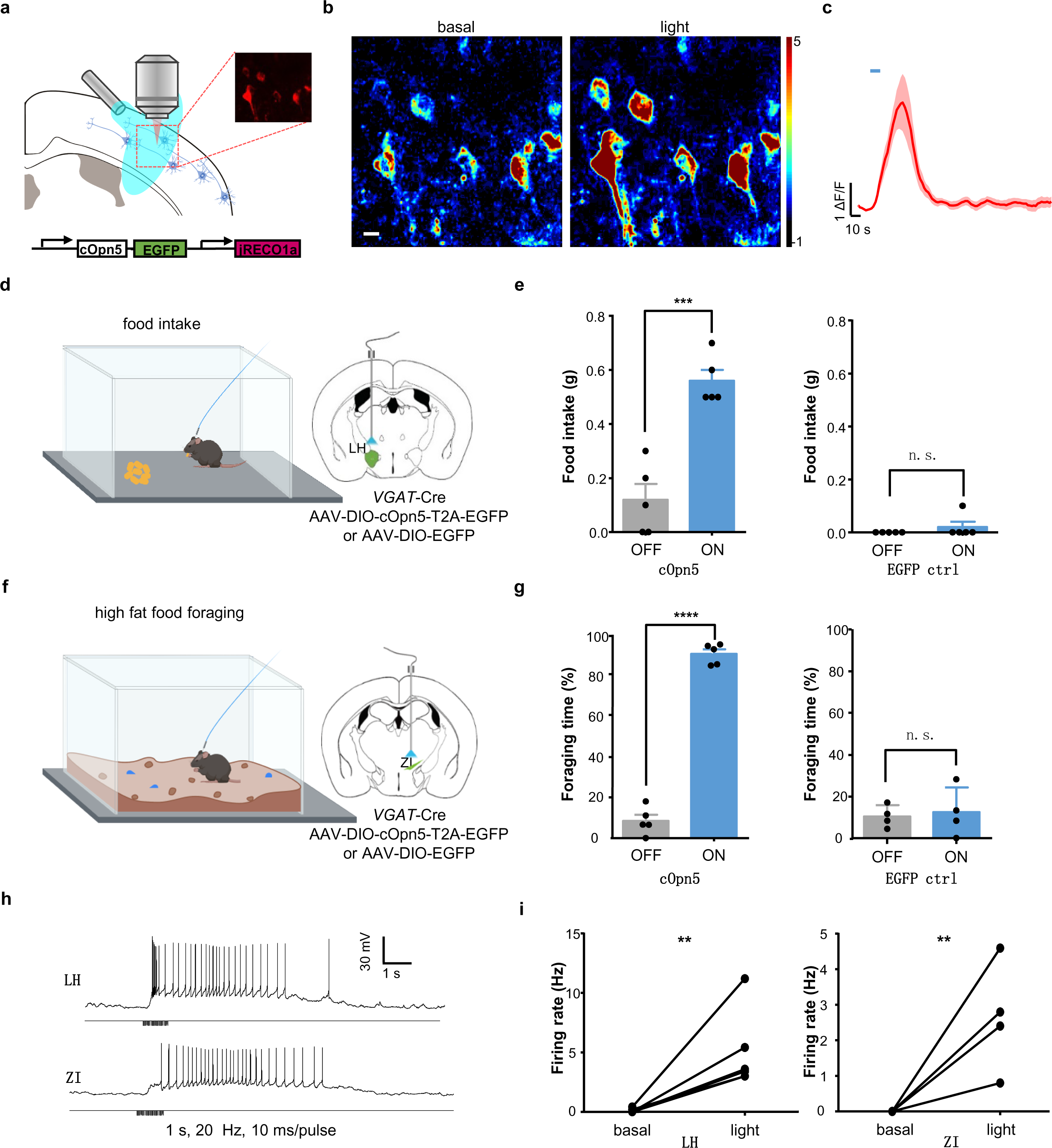
cOpn5-mediated optogenetics activation of neurons and changes of mouse behaviors in a neural circuit-dependent manner. **a,** Schematic diagram shows the experimental setup for optogenetic stimulation and Ca^2+^ imaging. We used AAV vectors to express cOpn5, EGFP, and the red Ca^2+^ sensor jRECO1a in neurons. **b,** Pseudocolor images show Ca^2+^ signals before and after light stimulation (10 s; 100 μW/mm^2^; 473 nm). Scale bar, 10 μm. **c**, Group data of Ca^2+^ signal traces of 6 individual neurons shown in **d.**, Schematic diagram of the experimental setup for cOpn5 optogenetics and food intake assay. cOpn5 and EGFP were expressed in GABAergic neurons within the LH of *VGAT*-Cre mice. EGFP was expressed as a control. **e**, Summary data show the light-induced (20 Hz; 5 ms/pulse; 473 nm; 0.75 mW output from the fiber tip), cOpn5-mediated activation of eating behaviors. ***P = 0.0003; n.s., non-significant; N = 6 mice; unpaired *t* test. Error bars indicate S.E.M.. **f**, Schematic diagram of the experimental setup for food foraging behavior. High-fat food pellets were used. cOpn5 and EGFP were expressed in GABAergic neurons within the ZI of *VGAT*-Cre mice. EGFP was expressed as a control. **g**, Summary data show the cOpn5-mediaed food foraging behaviors. Foraging time percentage was calculated upon receiving light stimulation until the mouse found the hidden food. ****P <0.0001; N = 6 mice; unpaired *t* test. Error bars indicate S.E.M.. **h**, Raw data illustrate the pattern of cOpn5-mediated optical activation of GABAergic neurons in these two brain areas. **i**, Group data show the neuronal firing rates before and after pulsed 473 nm light stimulation (1 Hz, 5 s; for the LH, N = 6 neurons, **P = 0.0041, unpaired t tests; for the ZI, N = 5 neurons, **P = 0.0027, unpaired t tests).

We next assessed the utility of cOpn5-mediated optogenetics for modulating animal behavior. The lateral hypothalamus (LH) participate in reward processing and feeding regulation^53–55^. We expressed cOpn5 in the LH GABAergic neurons of *VGAT-*Cre mice and implanted optical fibers to deliver light pulses into the LH of freely behaving mice (Fig. 3d and Extended Data Fig. 3b). Consistent with a role of LH GABA neurons in promoting feeding behavior^55^, light stimulation (20 Hz; 5 ms/pulse; 473 nm; 0.75 mW output from the fiber tip) elicited a significant increase in food intake in cOpn5-expressing mice but not the EGFP-expressing control mice (Fig. 3e). We also used a food-foraging behavior task to test the effect of cOpn5-mediated optogenetic activation of GABA neurons in the zona incerta (ZI) (Fig. 3f and Extended data Fig. 3c), a region known to drive compulsive eating^56^. cOpn5-expressing mice, but not the EGFP-expressing mice, showed a significantly increase in the time of foraging high fat food pellets upon repeated light stimulation (Fig. 3g). Notably, mice maintained the behavior (feeding behavior or high-fat food foraging behavior) while the light was on, and immediately stopped the behavior when the light was off (Supplementary Video 3). Thus, cOpn5 is effective for rapidly, accurately, and reversibly modulating animal behavioral states.

Finally, we investigated the effect of light-induced cOpn5 activation on the electrophysiological properties of lateral hypothalamus and zona incerta neurons in slice preparations (Extended Data Fig. 3a). Light pulses rapidly drove vigorous firing of action potentials (Figs. 3h and 3i). What’s more, we observed two types of activation patterns of cortical and subcortical neurons. In a majority of neurons (17 out 29 neurons), brief light pulses (10 ms) rapidly evoked strong inward currents (100-1000 pA) and drove vigorous firing of action potentials (Extended Data Fig. 3b, left and 3c). In the other 12 out of the 29 neurons recorded, blue light pulses (10 ms) induced a small depolarizing current (∼20 pA) in the voltage-clamp mode, and induced delayed-yet-robust firing of action potentials in the current-clamp mode (Extended Data Fig. 3b, right and 3c). Neurons were repeatedly stimulated with 10 ms pulses at 10 Hz, and exhibited a non-attenuated mode in firing rate across repetitive trials of light stimulation (Extended Data Fig. 3d). Of note, unlike those generated by ChR2 optogenetics^57^, the action potentials evoked by cOpn5 photostimulation were not time-locked to light pulses in any of neurons recorded.

### cOpn5 optogenetic activation of astrocytes evokes calcium signal

We extended the findings into primary cell cultures. It was reported transient elevations of calcium concentration occur in astrocytes providing cells with a specific form of excitability^58–61^. We expressed cOpn5 in primary astrocyte cultures prepared from the neonatal mouse brain with AAV vectors for bicistronic expression of cOpn5 and the EGFP marker protein (Fig. 4c). Using the Calbryte^TM^ 630 AM dye to monitor Ca^2+^ levels, we found that blue light illumination of cOpn5-expressing astrocytes produced strong Ca^2+^ transients (∼8 *ΔF/F*) (Figs. 4a, 4b and Supplementary Video 1). Confining the light pulses (63 ms) to subcellular regions produced Ca^2+^ signal propagation within the stimulated cells (Fig. 4d). Resembling the tests in HEK 293T cells, we observed wave-like propagation of Ca^2+^ signals that proceeded from the stimulated astrocyte gradually to more distal, non-stimulated, astrocytes (Figs. 4e, 4f and Supplementary Video 2). This optically-triggered intercellular Ca^2+^ wave was reminiscent of the dynamics of astrocytic networks that were initially discovered using neurochemical and mechanical stimulation^58, 59^.

**Fig. 4.**
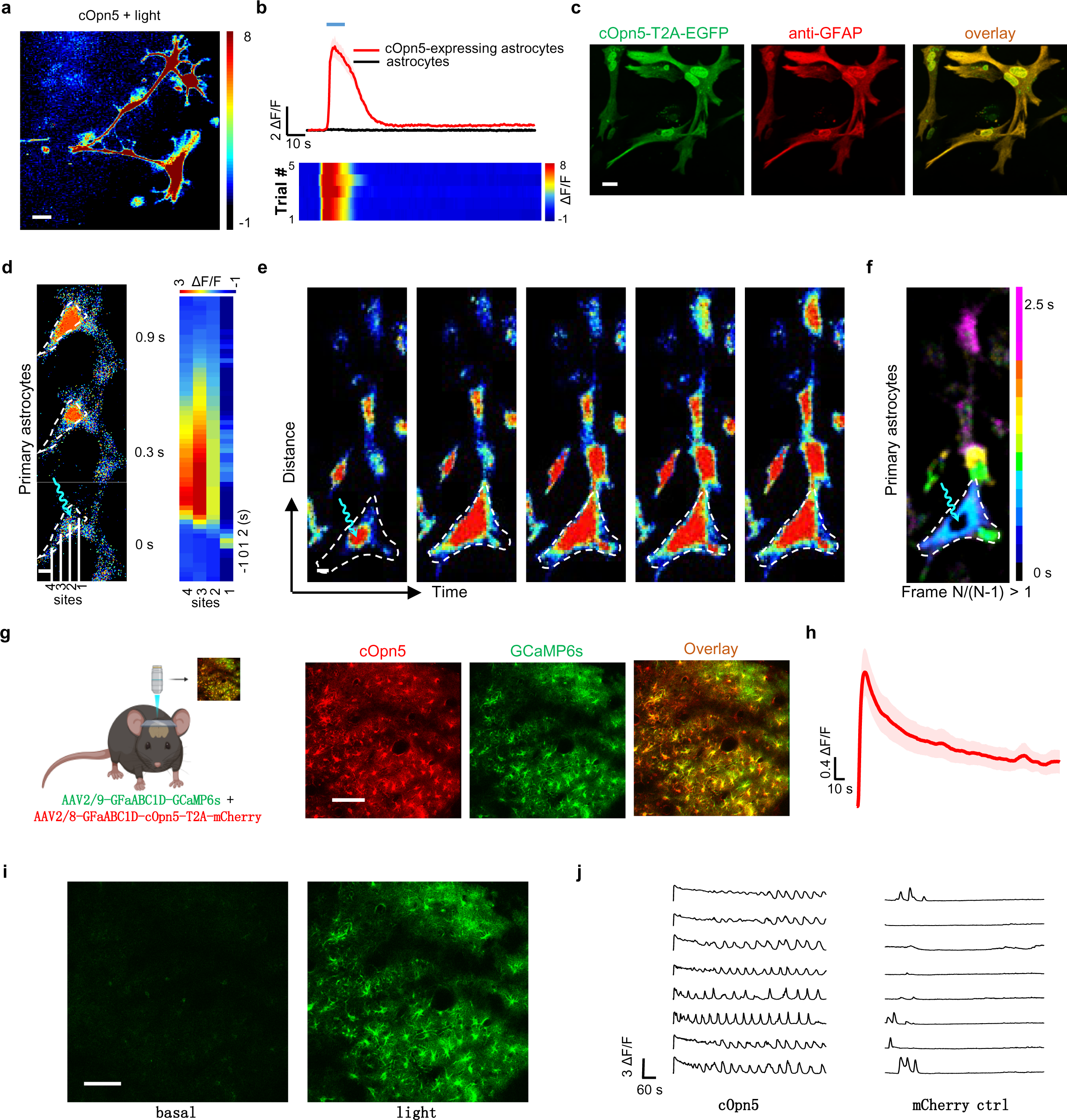
cOpn5-mediated optical activation of astrocytes evokes calcium signal. **a**, Pseudocolor images of the baseline and peak Ca^2+^ signals following light stimulation of cOpn5-expressing astrocytes. Scale bar, 20 μm. **b**, Plot of Ca^2+^ signals and heat map representation of Ca^2+^ signals across trials (n = 25 astrocytes). **c**, cOpn5 was expressed in cultured primary astrocytes using AAV-cOpn5-T2A-EGFP (green). Astrocyte identity was confirmed by GFAP immunostaining (red). Scale bar, 20 μm. **d**, Images of light-induced Ca^2+^ signal propagation in a single cOpn5-expressing primary astrocyte stimulated in a subcellular region (stimulation size 4×4 μm^2^ and frame interval = 300 ms). Scale bar, 10 μm. **e**, Images of light-induced Ca^2+^ signal propagation in cOpn5-expressing primary astrocytes. Scale bar, 10 μm. **f**, Pseudocolor images showing the process of Ca^2+^ signal propagation across time of **g** (frame N/(N-1) > 1). Frame interval was 500 ms and each frame is counted once. **g**, Schematic diagram of the experimental setup for *in vivo* two-photon imaging (920 nm) of astrocyte Ca^2+^ imaging following cOpn5-mediated astrocyte activation. Images show the expression of GCaMP6s (green) and cOpn5-T2A-mCherry (red) in astrocytes within the mouse S1 cortex. Scale bar, 100 μm. **h**, Time courses of light-evoked Ca^2+^ signals in astrocytes (N = 5). **i**, Images of the Ca^2+^ signals before (basal) and 5s after light stimulation (light). Scale bar, 100 μm. **j**, Raw traces of eight individual GCaMP6s-expressing astrocytes signals in a mouse that expressed cOpn5 or mCherry.

We next tested the performance of cOpn5-mediated optogenetics *in vivo*. We carried out cOpn5-mediated optogenetic activation of astrocytes and monitored calcium signal^60^. Specifically, we expressed cOpn5 and the GCaMP6s sensor in the mouse S1 sensory cortex following the infusion of AAV vectors containing the GfaABC1D promoter. (Fig. 4g), which drove gene expression selectively and efficiently in cortical astrocytes^62, 63^. We initially expected that, in addition to the 920 nm light from pulsed laser for two-photon imaging, blue light pulses would be required to stimulate calcium signals. Strikingly, upon the 920 nm light delivered from microscope, triggered remarkable calcium elevation was observed in the cOpn5- and GCaMP6s-expressing mice but not in the control mice that expressed the mCherry- and GCaMP6s but lacked cOpn5 expression (Figs. 4h-4j and Supplementary Video 4). The calcium events were persistent with only slight decrease during 20min constant stimulation (Extended Data Fig. 4a). Together, these experiments demonstrate that cOpn5 optogenetics allows precise spatiotemporal control of G_q_ signaling with millisecond and subcellular resolutions.

### cOpn5 optogenetic activation of astrocytes induces massive ATP release and regulates neuron activation *in vivo*

Astrocytes represent an important population of non-excitable cells in the central nervous system and are known to regulate a variety of processes, including neurogenesis and synaptogenesis, blood-brain barrier permeability, and extracellular homeostasis^64–68^. However, to date direct optogenetic control of astrocytes has achieved only limited success^28, 69, 70^. ATP is known as a key messenger for inter-astrocyte communication^71–73^. In cell cultures and brain slices, ATP can be released through hemichannels in a Ca^2+^-independent manner or through vesicle release in a Ca^2+^-dependent manner^74, 75^. It has remained controversial whether intracellular Ca^2+^ signals in astrocyte induce ATP release *in vivo*, and if so, how such release manifests in real-time. We carried out cOpn5-mediated optogenetic activation of astrocytes and monitored ATP release using the recently-developed GPCR Activation‒Based ATP sensor GRAB_ATP_^76, 77^. Specifically, we expressed cOpn5 and the GRAB_ATP_ sensor in the mouse S1 sensory cortex following the infusion of AAV vectors containing the GfaABC1D promoter (Fig. 5a).We tested the gene was selectively and efficiently expressed in cortical astrocytes (Figs. 5b and 5c)^62, 63^. We also used 920 nm light from pulsed laser for two-photon imaging. The 920 nm light could also triggered massive ATP flashes in the cOpn5- and GRAB_ATP_-expressing mice, but not in mice that expressed the ATP sensor but lacked cOpn5 expression (Fig. 5e). Individual ATP flashes typically ranged in diameters of 20-100 μm and lasted for ∼1 min (Supplementary Video 5). The flash frequency gradually increased following ∼1 min of initial quiescence and up to the level of ∼50 flashes per min within the imaging area (636×636 μm^2^) in ∼5 min (Figs. 5d and 5f). In mice expressing GRAB_ATP_ alone, we observed sporadic ATP events (∼0.3 flashes per min within the imaging area; Figs. 5e and 5g). The observed ATP flash frequency in cOpn5-expressing mice was ∼1300 times more than non-cOpn5-expressing mice (Figs. 5e, 5g and Supplementary Video 5). Moreover, high-frequency ATP flashes also occurred in the repeated trials (Extended Data Fig. 4b). Given that hM3Dq allows the activation of astrocytes with Ca^2+^ elevation by CNO^60, 78^, we also performed ATP imaging 40 min following the treatment of intraperitoneal injections of CNO into GRAB_ATP_-hM3Dq expressing mice that lacked cOpn5 expression. Following the CNO treatments, the ATP flash events were approximately equal to the basal condition (Figs. 5e and 5g and Supplementary Video 5) and hM3Dq was ineffectively to trigger astrocytes activation-dependent ATP release. These experiments thus demonstrate that cOpn5-mediated optogenetic activation of G_q_ signaling in astrocytes is particularly effective in inducing ATP release *in vivo* in the unique form of ATP flashes.

**Fig. 5:**
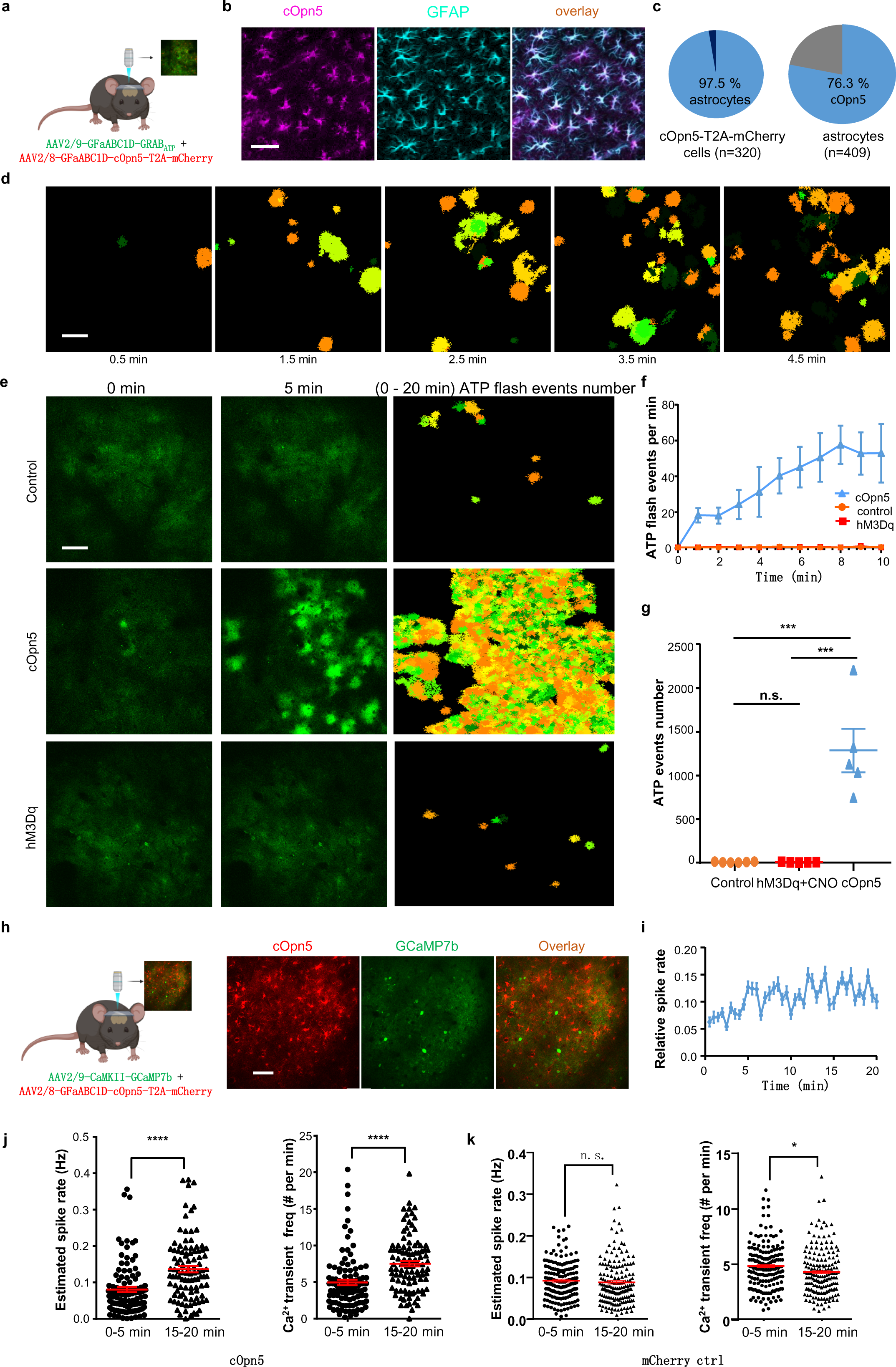
cOpn5-mediated optogenetic activation of astrocytes induces massive ATP flashes and neuronal activation *in vivo*. **a**, Schematic diagram of the experimental setup for *in vivo* two-photon imaging (920 nm) of ATP signal following cOpn5-mediated optical activation of astrocytes. We infused two AAV vectors to express cOpn5-T2A-mCherry (red), and GRAB_ATP_ sensor (green) in the astrocytes within the mouse S1 cortex. **b**, Pseudocolor images show that AAV-GfaABC1D-cOpn5-T2A-mCherry drove mCherry expression in S1 cortical cells that were immunopositive to GFAP, a marker of astrocytes. Scale bar, 20 μm. **c**, Out of 320 cOpn5-mCherry+ cells examined, 312 cells were GFAP+, indicating 97.5% precision. Within the same imaged areas, a total of 409 cells were GFAP, indicating 76.3% efficiency of astrocyte labeling using the AAV-GFaABC1D constructs. **d**, Representative images show astrocytic ATP flash events in a cOpn5-expressing mouse in the first 4.5min, with different colors indicating individual flashes. Scale bar, 100 μm. **e**, Images show the overall ATP flash events in a control mouse (no cOpn5 expression), a CNO (2 mg/kg) - treated hM3Dq expression mouse (no cOpn5 expression) and a cOpn5-expressing mouse. The left column shows the raw GRAB_ATP_ images at the basal level (before light delivery); the middle column shows GRAB_ATP_ signals at 5 min; and the right column shows pseudo-color-coded ATP flash events accumulated during 0-20 min. Scale bar, 100 μm. **f**, The plot of astrocytic ATP flash events number per min across time (0-10 min) in control group (no cOpn5 expression), CNO-treated hM3Dq expression group (no cOpn5 expression), and cOpn5-expressing group. **g**, The quantification of ATP flash events number in 20min of the control (N = 6), hM3Dq+CNO (N = 5), and cOpn5 groups (N = 5). For the control - hM3Dq+CNO comparison, no significant difference; for the control - cOpn5 comparison, ***P = 0.0003, for the hM3Dq+CNO - cOpn5 comparison, ***P = 0.0009; unpaired *t* tests. **h**, Schematic diagram of the experimental setup for *in vivo* two-photon imaging (920 nm) of neuron Ca^2+^ imaging following cOpn5-mediated astrocyte activation. Images show the expression of cOpn5-T2A-mCherry (red) in astrocytes and the expression of a GCaMP7b (green) in neurons within the mouse S1 cortex. Scale bar, 100 μm. **i**, The plot of average spike rate relative to basal across time (0-20 min). N=193 neurons from 3 mice **j**, Decoded spike rate analysis of GCaMP7b-expressing neurons within the periods of 0-5 min and 15-20 min that were coupled with cOpn5-mediated astrocytes activation (left panel) N = 193 neurons). ****P < 0.0001, unpaired *t* test. The number of Ca^2+^ transients of GCaMP7b-expressing neurons during 0-5 min and 15-20 min, coupled with cOpn5-mediated astrocytes activation (right panel). N=193 neurons, ****P < 0.0001, Unpaired *t* test. **k**, Decoded spike rate analysis of GCaMP7b-expressing neurons within the periods of 0-5 min and 15-20 min that were coupled with imaging from mice with mCherry expression in astrocytes as control (left panel) (N = 170 neurons). no significant difference; unpaired *t* test. The number of Ca^2+^ transient analysis of GCaMP7b-expressing neurons during the periods of 0-5 min and 15-20 min in mice that expressed mCherry in astrocytes (right panel). N = 170 neurons, *P =0.0146, paired *t* test.

Astrocytes release several gliotransmitters, which together with their metabolites, exert complex modulatory effects on synaptic activity, neuronal oscillation, and various animal behaviors^79, 80^. For example, the gliotransmitters ATP and glutamate can directly activate nearby neurons^64, 72, 81, 82^, whereas adenosine, the major metabolite of ATP, may inhibit neurons through the A1-type adenosine receptor^83^. It has remained elusive how astrocyte activation affects the activity of individual neurons in behaving animals. We expressed cOpn5 in astrocytes and the green Ca^2+^ indicator GCaMP7 in pyramidal neurons within the S1 cortical area. We then carried out two-photon imaging of neuronal Ca^2+^ signals in response to cOpn5-mediated optical activation of astrocytes in head-fixed awake, behaving mice (Fig. 5h). Initially neurons were largely quiescent, and then exhibited a gradual increase in the frequency and amplitudes of Ca^2+^ transients in approximately 5 min (Fig. 5i). Group data from a total of 193 neurons revealed that photostimulation of cOpn5-expressing significantly increased the frequency of Ca^2+^ transients and the in neurons. MCherry-expressing control group showed no influence on the decoded spiking rates of neurons or even slightly decrease the Ca^2+^ transients frequency (Figs. 5i, 5j and Extended Data Figs. 4c, 4d), suggesting that activating G_q_ signaling in astrocytes produces an overall excitatory effect on nearby cortical neurons in behaving state. Both astrocytic ATP imaging and neuronal Ca^2+^ imaging demonstrated that the long-wavelength (920 nm) light from pulsed laser for two-photon imaging is able to activate cOpn5, indicating the possibility of two-photon optogenetics for cOpn5.

## Discussion

Here, we demonstrate the use of cOpn5 as an extremely effective optogenetic tool for activating G_q_ signaling. Previous studies have characterized mammalian Opn5 as a UV-sensitive G_i_-coupled opsin; we present the surprising finding that visible blue light can induce rapid Ca^2+^ transients, IP_1_ accumulation, and PKC activation in cOpn5-expressing mammalian cells. cOpn5 in mouse astrocytes effectively mediates light-evoked ATP release and elevates neuron activity *in vivo*. We also show that cOpn5 allows optical activation of neurons and precise control of animal behaviors. Importantly, cOpn5 is a powerful yet easy-to-use, single-component system that does not require an exogenous chromophore. We envision that cOpn5-based optogenetics will be an enabling technique for investigating the important physiological and behavioral functions regulated by G_q_ signaling in both non-excitable and excitable cells.

Supplementary Table 4 lists the key features of cOpn5 by directly comparing its response amplitudes, light sensitivity, temporal resolution, and the requirement of additional chromophores to those of other optogenetic tools. For cOpn5-expressing cells, merely 10 ms blue light pulses at the intensity of 16 μW/mm^2^ evoke rapid increase in Ca^2+^ signals with the peak amplitudes of 3-8 *ΔF/F*. By contrast, prior characterizations and this study show that the activation of opto-a1AR or mammalian Opn4, the two proposed optogenetic tools for G_q_ signaling, require ∼3-order higher light intensity (7-40 mW/mm^2^) and prolonged light exposure (20-60 s) but produce only weak Ca^2+^ signals (0.25-1 *ΔF/F*). Therefore, opto-a1AR or mammalian Opn4 cannot mimic the rapid activation profiles of endogenous G_q_-coupled receptors that often trigger strong Ca^2+^ release upon the application of their corresponding ligands. We demonstrate the power of cOpn5 optogenetics by showing the striking physiological and behavioral effects in response to cOpn5-mediated optical activation of astrocytes and neurons *in vivo*. By contrast, recent *ex vivo* studies show that opto-a1AR- and Opn4-mediated optogenetic stimulations slightly increase the amplitudes of Ca^2+^ signals and only mildly modulate the frequency of Ca^2+^ transients and synaptic events even after prolonged illumination^28, 29^. By overcoming the limitations of light sensitivity, temporal resolution, and response amplitudes associated with opto-a1AR- and Opn4-mediated optogenetics, cOpn5 should find broad applicability for studying G_q_ signaling in numerous cells and tissues.

cOpn5 optogenetics also enjoys the benefits of safety and convenience. Although Opn5 from many species are reported UV-responsive^36^, cOpn5 is optimally activated by 470 nm blue light, which penetrates better than UV and avoids UV-associated cellular toxicity. Its ultra-sensitivity to light minimizes potential heating artifact. It is two-photon activable using long-wavelength light (920 nm here), suggesting that it is suitable for even deeper tissue activation using a pulsed laser. cOpn5 is strongly, and repetitively activated by light without the requirement for exogenous chromophore, possibly because cOpn5 is a bistable opsin that covalently binds to endogenous retinal and is thus resistant to photobleaching^32, 84^. By contrast, some studies indicate that the mammalian Opn4 requires the addition of chromophore for continuous activation^25, 26^. cOpn5 as a single-component system is particularly useful for both *in vitro* and *in vivo* studies as it avoids the burden of delivering a compound into the tissue during the experiment.

cOpn5 optogenetics has several major advantages over chemogenetics and uncaging tools. It is temporally much more precise and offers single-cell or even subcellular spatial resolution. Although CNO/hM3Dq-mediated chemogenetics has been used to investigate the physiological and behavioral functions of non-excitable cells, such as astrocytes in the brain^85, 86^. The diffusive nature of compounds indicates that it is nearly impossible to chemogenetically stimulate G_q_ signaling with cellular and subcellular resolution. cOpn5 also differs from caged compound-based ‘uncaging’ tools such as caged calcium and caged IP_3_, since these tools require compound preloading and only partially mimic the Ca^2+^-related pathways associated with G_q_ signaling. There exist other ‘uncaging’ tools, such as caged glutamate and caged ATP^87, 88^, that target endogenous receptors. However, these caged compounds lack cell selectivity and require their introduction into extracellular medium or the intracellular cytoplasm, which limits their applications in behaving animals.

cOpn5 optogenetics should be particularly useful for precisely activating intracellular G_q_ signaling, which subsequently triggers Ca^2+^ release from intracellular stores and activates PKC. cOpn5 differs from current channel-based optogenetic tools, such as ChR2 and its variants, which translocate cations across the plasma membrane. By controlling cellular membrane potentials and thus action potential firing, ChR2 and its variants have contributed tremendously to functional dissection of neural circuits; however, their successes have been more constrained in studying non-excitable cells that lack active ion channels for generating action potentials^13^. In addition to the applications on non-excitable cells, cOpn5 optogenetics can also stimulate G_q_ signaling in neurons and control animal behavior in a circuit-dependent manner. Of note, G_q_-coupled GPCRs may affect variable downstream signaling in a receptor- and cell-specific manner^89^. Indeed, we observed that the same light illumination parameters produce different activation patterns among neurons. We recommend characterizing and confirming the activation profiles, similar to the applications of other optogenetic and chemogenetic tools. cOpn5-mediated optogenetic activation does not generate strictly time-locked action potential firing as precisely as that by ChR2 in neurons. Ion channel-based optogenetic tools would be preferable if temporally precise control of action potential firing is necessary. Nevertheless, cOpn5-mediated asynchronous firing activity may be useful for circuit dissection, since it avoids the potential artefact of massively synchronized neuronal activation.

In addition to the technical advances, our findings also have several functional implications about astrocyte functions *in vivo*. Although ATP is considered an important gliotransmitter, previous studies have revealed multiple releasing mechanisms that depend on the methods of stimulation (electrical, neurochemical, or mechanical), the presence of extracellular Ca^2+^, and the exact form of cell and tissue preparations^30, 90^. Though it was reported astrocytes activation could evoke an ATP/Adenosine-dependent transient boost^28^, the presence of ATP release is monitored indirectly. It remained controversial whether activating G_q_ signals in astrocytes triggers ATP release, and if so, how this release is expressed *in vivo*. Here we provide the first demonstration that stimulating G_q_ signaling pathway within astrocytes triggers massive ATP release in the form of ATP flashes. cOpn5 optogenetics thus provides an ideal technique to study the molecular and cellular mechanisms underlying ATP release. In addition to ATP, astrocyte activation leads to the release of other gliotransmitters, such as D-serine, glutamate, and GABA^81^. The gliotransmitters and their metabolites can exert complex modulatory effects on neuronal excitability and synaptic strength^88, 91^. It had remained unclear how these various effects are integrated to modulate neuronal activity *in vivo*. Here we reveal that optogenetic activation of cOpn5-expressing astrocytes leads to an overall excitatory effect on pyramidal neurons in the S1 cortex of mice. This optogenetic approach lays the foundation for dissecting the molecular, cellular, and circuit mechanisms underlying the rich interactions between astrocytes and neurons. Our experiments *in vivo* have also demonstrated that cOpn5 has compatibility with optogenetic probes and imaging sensors, such as genetically encoded Ca^2+^ sensors and GPCR-based neurotransmitter sensors^57, 92–96^. cOpn5 together with these sensors potentially allows an all-optical approach to activate G_q_ signaling and simultaneously monitor the relevant effects^97^.

In summary, we present cOpn5 as a blue light-sensitive opsin for rapidly, reversibly, and precisely activating G_q_ signaling. We also establish cOpn5 as a powerful and easy-to-use optogenetic tool for activating both non-excitable cells and neurons. Given the ubiquitously important roles of G_q_-coupled GPCRs, we expect that cOpn5 will find broad applications for studying the mechanisms and functions of G_q_ signaling in all major cell types and tissues.

---

**Extended Data Fig. 1.**
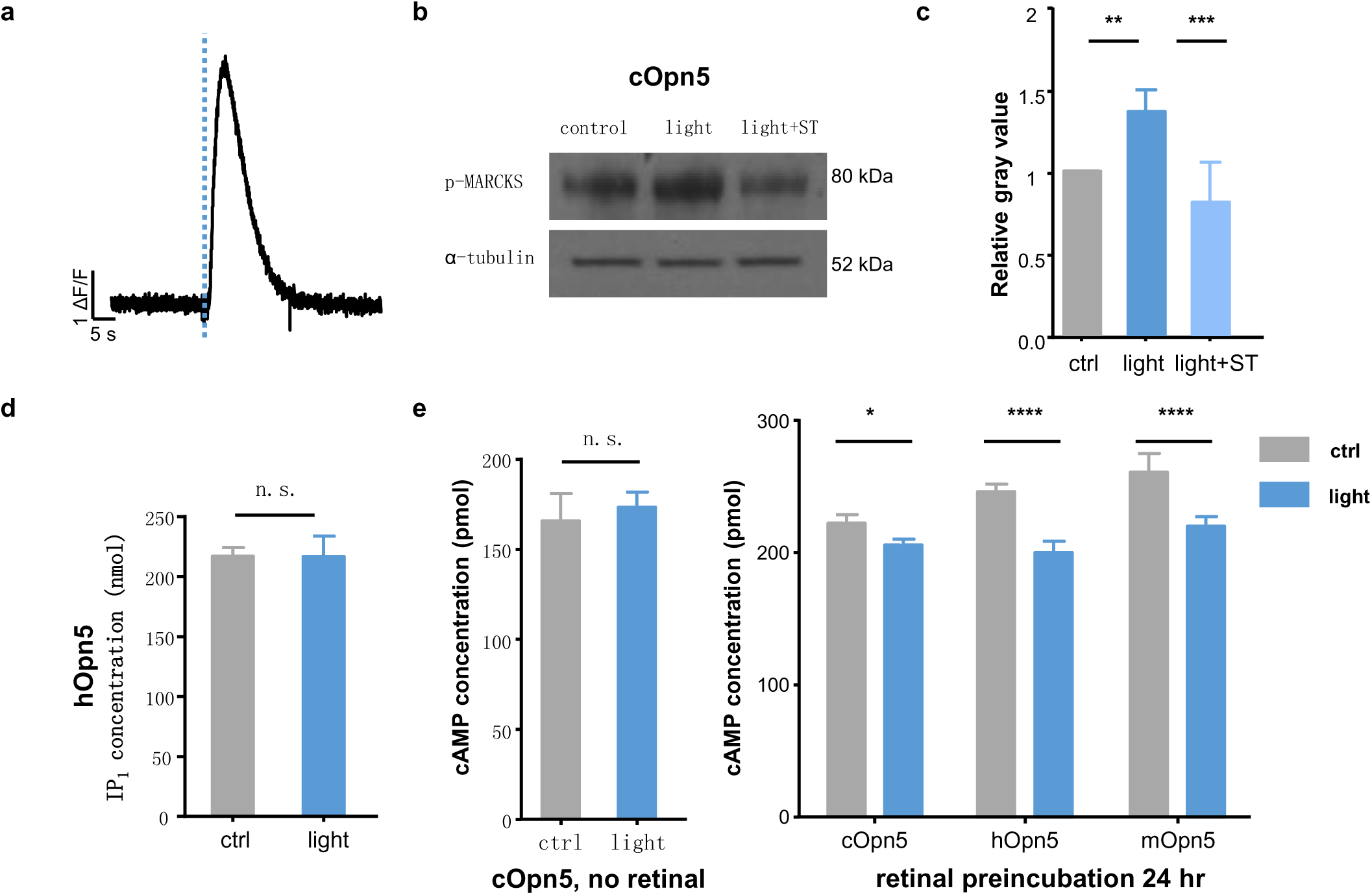
cOpn5 couples to G_q_ signaling. **a**, Time course of optically evoked Ca^2+^ signal in extracellular Ca^2+^ free medium (10 ms; 16 μW/mm^2^; 470 nm; N = 17 HEK 293T cells). **b**, One representative of phosphorylation of MARCKS in cOpn5-expressing HEK 293T cells in the control group (no light stimulation), the light stimulation group, and light+staurosporine (ST) group (ST, a PKC inhibitor; 10 μM) without addition of retinal. **c**, The amount of p-MARCKS was normalized to the amount of α-tubulin in the same fraction. N = 4, **P = 0.0096, ***P = 0.0004; Tukey‘s multiple comparisons test. **d**, IP_1_ accumulation in human Opn5-expressing HEK 293T cells with or without light stimulation without addition of retinal. N = 4, n.s., no significant difference; unpaired *t* test. **e**, Light has no effect on cAMP levels (10 μM forskolin preincubation) in cOpn5-expressing HEK 293T cells without additional retinal in the medium (left panel) (N = 4). Right panel shows the effects of photostimulation on cAMP concentrations for HEK 293T cells expressing Opn5s from four different species following 10 μM retinal Preincubation (N = 4). n.s., no significant difference; unpaired *t* test. Error bars in **c, d** and **e** indicate S.E.M..

**Extended Data Fig. 2.**
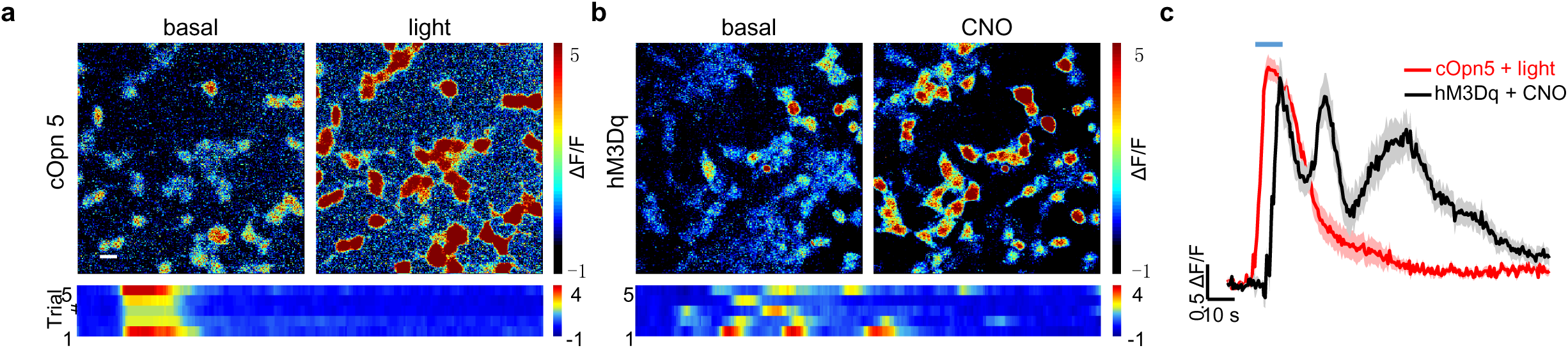
Direct comparisons between the performance of cOpn5 and hM3Dq chemogenetics. **a** Pseudocolor images (upper) and heat map representation of Ca^2+^ signals evoked by 10 s optical stimulation (same as the CNO stimulation time) of cOpn5-expressing cells across 5 consecutive trials. Scale bar, 20 μm. **b,** Effect of chemogenetic stimulation on the Ca^2+^ signals in hM3Dq-expressing HEK 293T cells. **c**, Time courses of Ca^2+^ signals evoked by cOpn5-mediated optogenetic stimulation (10 s) and hM3Dq-mediated chemogenetic stimulation using CNO puff (100 nM; 10 s), respectively (N = 20, cOpn5-expressing HEK 293T cells and 20 hM3Dq-expressing HEK 293T cells).

**Extended Data Fig. 3.**
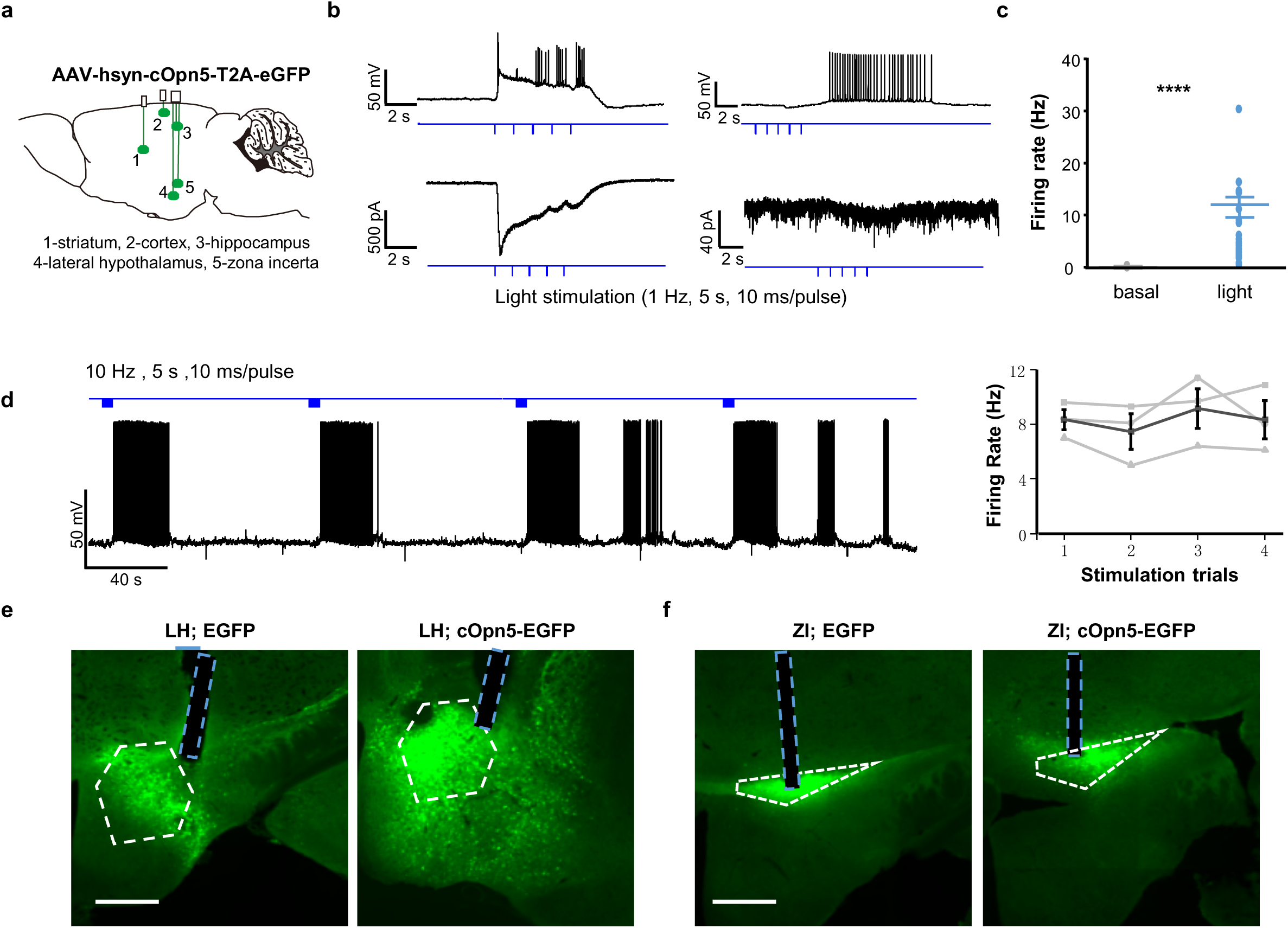
cOpn5-mediated optogenetics reliably activates neurons. **a**, Schematic diagram depicts optogenetic stimulation and whole-cell patch-clamp recording of cOpn5-expressing neurons in the cortex, striatum, hippocampus, lateral hypothalamus (LH), and zonal incerta (ZI). **b**, Raw data illustrate two representative patterns of light-evoked neuronal activation. One neuron (left panels) exhibited rapid membrane potential depolarization and large inward currents (1 Hz, 5s, 10 ms/pulse), and another neuron exhibited strong, delayed firing of action potentials yet small sustained inward currents in response to the light pulses. **c**, Group data show the neuronal firing rates before and after pulsed 473 nm light stimulation (20 Hz, 1 s; N = 29 neurons; ****P < 0.0001, unpaired *t* test). **d**, Raw trace shows that cOpn5 mediated reliable and reproducible photoactivation of a neuron (left). The right panel shows the summary of firing rates across repetitive trials of light stimulation (N = 3 neurons). **e**, Images show the expression of EGFP control and bicistronic expression of EGFP and cOpn5 in the LH (white dashed lines). Lesion sites and blue dashed lines indicate the placement of optical fibers. Scale bars, 500 μm. **f**, The injection sites and optical fiber placement in the ZI. Scale bars, 500 μm.

**Extended Data Fig. 4.**
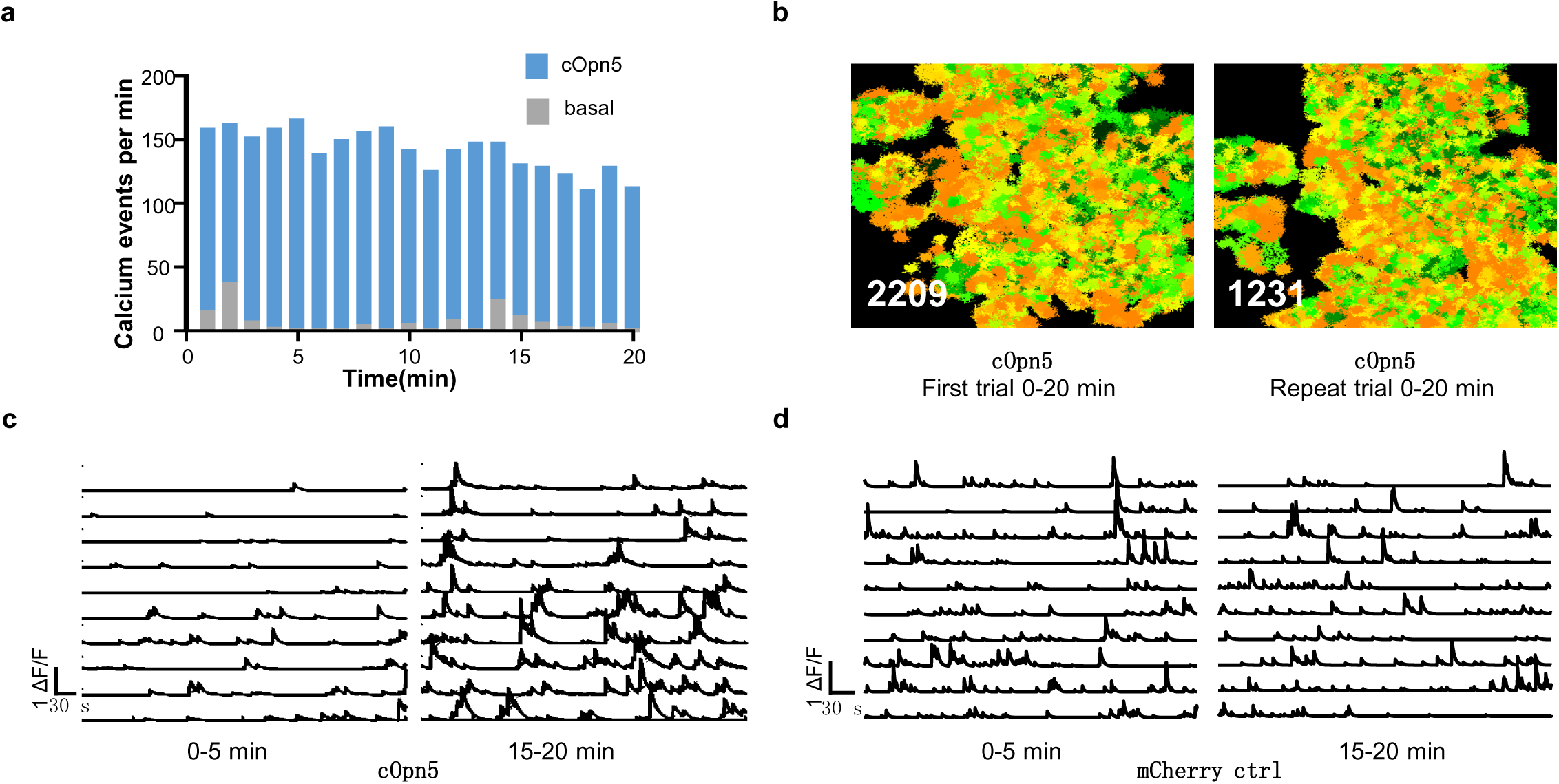
cOpn5-mediated optical activation of astrocytes, produces reliable ATP release and the activation of surrounding neurons. **a**, Time courses of light-evoked Ca^2+^ events in astrocytes for 20min constant stimulation (N = 5). **b**, Pseudocolor color image show total ATP flash events (0-20 min) in cOpn5-expressing mouse in a repeat trial 1 hour after the initial trial. **c**, Raw traces of ten individual GCaMP7b-expressing neurons signals in 0-5 min and 15-20 min, coupled with cOpn5-mediated optical activation of astrocytes . **d**, Raw traces of ten individual GCaMP7b-expressing neurons signals during the periods of 0-5 min and 15-20 min in a mouse that expressed mCherry in astrocytes.

**Supplementary Table 1: Comparison of chicken, human, and mouse Opn5.**

**Supplementary Table 2: Key resources.**

**Supplementary Table 3: Summary of statistical analyses.**

**Supplementary Table 4: Comparison cOpn5 with other optogenetic and chemogenetic tools.**

**Video 1.** cOpn5-mediated optical activation of Ca^2+^ signals in HEK 293T cells (left) and astrocytes (right). Related to Figure 1 and Figure 4. Caption indicated the period of the light on. Scale bars, 500 μm.

**Video 2.** Subcellular stimulation and Ca^2+^ waves in HEK 293 cells (left) and in astrocytes (right). Related to Figure 2 and Figure 4. Caption indicated the restricting brief light stimulation (63 ms) and the blue dotted arrows indicated the stimulation sites.

**Video 3.** cOpn5-mediated optogenetics changes mouse behaviors. The video first shows the behavior effect of optogenetically activating ZI GABAergic neurons and then the behavior effect of activating LH GABAergic neurons. Related to Figure 3.

**Video 4.** cOpn5-mediated optical activation of Ca^2+^ signals in a control mouse and cOpn5-expressing mouse with blue light stimulation. Related to Figure 4.

**Video 5.** ATP flash events in a control mouse (no cOpn5 expression), a mouse 40min after CNO treatment (hM3Dq expression), a cOpn5-expressing mouse during the initial imaging period (lower left) and during the second imaging session (lower right). Related to Figure 5.

