## Supplementary material for "A Neuropsin-based Optogenetic Tool for Precise Control of G_q_ signaling": methods

**Mice**

Animal care and use conformed to the institutional guidelines of the National Institute of Biological Sciences, Beijing as well as the governmental regulations of China. Male C57BL/6N mice were purchased from Beijing Vital River Laboratory Animal Technology Co., Ltd (China). Adult (8-16 weeks old) *VGAT*-Cre mice [STOCK Tg(*Slc32a1*-cre)2.1Hzo/FrkJ] of either sex were obtained from the Jackson Laboratory (USA). All mice were maintained with a 12/12 light/dark cycle (lights off at 8PM) and housed in groups of five for 6-8 weeks. After surgery, mice were housed in groups with an inverted light dark cycle (lights off at 8AM) for at least one week before further experiments. Adult mice of either sex were used. We used simplified genotypes of mouse strains for clarity.

**HEK 293T cell culture**

HEK 293T cells were obtained from the American Type Culture Collection (ATCC). Cells were cultured at 37ºC in 5% $CO_2$ in DMEM supplemented with 10% (v/v) fetal bovine serum (FBS) and 1% penicillin-streptomycin. HEK 293T stable cell lines were generated using lentivirus; transduced monoclonal cells were selected to perform the experiments. Antibiotic selection was used to maintain long-term gene expression.

**Primary astrocyte culture**

Primary astrocyte cultures were prepared from the cerebral cortices of mouse pubs (P0) as described previously [1]. Briefly, cortices were dissected free of meninges and placed in DMEM/F12 containing 1% penicillin-streptomycin and no fetal bovine serum (FBS). Following mechanical dissociation and filtration with 40 μm cell sieves, the cells were collected by centrifugation (1000 rpm) at room temperature for 10 min. The cells were suspended in DMEM/F12 medium of 10% FBS and 1% penicillin-streptomycin and plated in the culture flask. Cultures were grown for 14 days in a 37 °C incubation chamber with 5% $CO_2$ with regular media changes. Following shaking on a 37° shaker for 2 hours at 180-200 rpm, we retrieved the cells left at the bottom of the flask.

### Method Details

**DNA plasmids and Viral constructs**

Chicken *Opn5*, human *Opn5*, mouse *Opn5* and mouse *Opn4L* sequences were synthesized (Azenta Life Science). Supplementary Table 1 provides a detailed list of the three opsins, their species, GenBank accession numbers and the aliases used in the main text of this paper. Cell-filling versions of all the opsins were constructed using a ribosomal skip site (T2A) between the opsin and fluorescent proteins. DNA fragments were generated using PCR amplification with primers containing 20 base-pair overlap (Azenta Life Science) and then assembled into plasmids using Gibson assembly (Supplementary Table 2). For cultured cell studies, all genes were subcloned into a lentiviral backbone pLJM1 vector (Addgene plasmid #19319) under a CMV promoter to make vector *pLJM1-Opn5*. *cOpn5* gene was subcloned into *pLJM1* vector with C-terminal V5 tag fusion and *T2A-EGFP* for bicistronic expression.

For *in vivo* neuron studies, cOpn5 gene was subcloned into pAAV vector under the human synapsin promoter (hSyn) promoter or the CaMKII promoter with T2A-EGFP bicistronic expression. *cOpn5* used in astrocytes was also subcloned into a pAAV backbone, but under a *GfaABC1D* promoter instead with mCherry fusion. The Cre-dependent *cOpn5-T2A-eGFP* vector was generated using the vector with lox sites flanking the protein-coding region under *EF1a* promoter. All restriction enzymes used for molecular cloning were from New England Biolabs and the constructs were verified using Sanger sequencing. The plasmids were amplified and then purified using Endo Free Plasmid Maxi kit (Vazyme, Cat. # DC202-01). The *pcDNA3.1/opto-a1AR-EYFP* plasmids were obtained through Addgene (plasmid #20947). *CAAX–EGFP* was a gift from Yulong Li (Peking University, China).

AAV vectors carrying *EF1a-DIO-cOpn5-T2A-eGFP* and *hSyn-cOpn5-T2A-eGFP* constructs were packaged into AAV2/9 serotype. *GfaABC1D-cOpn5-T2A-eGFP ,GfaABC1D-cOpn5-T2A-mCherry* were packaged into AAV2/8 serotype and *GfaABC1D-GCaMP6s* were packaged into AAV2/9 serotype with titers of 1-5×$10^{12}$ viral particles/ml (Vector Core at the Chinese Institute for Brain Research, Beijing). We purchased the *AAV2/9-GfaABC1D-ATP1.0* ($GRAB_{ATP}$ sensor) and *AAV9-hSyn-NES-jRGECO1a- WPRE* from WZ Biosciences Inc. with titers of 1×$10^{13}$ viral particles/ml. All Plasmids and Viral constructs used in this paper were listed in Supplementary Table 2.

**Common surgery and virus injection**

Adult mice were anesthetized with avertin (i.p. 300mg/kg) before surgery and then placed in a stereotaxic apparatus. After disinfection with 0.3% hydrogen peroxide, a small incision of the scalp was created to expose the skull. Craniotomy was conducted above the targeted brain areas using the following coordinates (mm in AP, DV, ML with respect to the bregma): −0.7, −4.75, ± 1.0 for the lateral hypothalamus, −1.0, −4.4, ± 0.7 for the zona incerta , -1.6, -0.7, -2.0 for the S1 cortex, 0.2, -3.5, 1.8 for the dorsal striatum, and 1.8, 1.9, 1.7 for the hippocampus. We used a microsyringe pump (Nanoliter 2010 Injector, WPI) and a Micro4 controller to slowly infused AAV virus into each site (46 nl/min speed; a total volume of 100 nl unless described otherwise).

**Immunohistochemistry, histology and fluorescent microscopy**

For immunofluorescence, cultured HEK 293T cells transiently-transfected with pLJM1-cmv-V5-Opn5 and primary astrocytes cells were washed with PBS for three times and fixed with 4% paraformaldehyde (PFA, wt/vol in PBS) for 20 min at room temperature (RT). The fixed cells were washed with PBS again and were then blocked with 3% FBS and 0.3% Triton X-100 for 30 min followed by staining with the primary antibodies for 2 hr at RT. Cells were washed with PBS for three times and stained with the secondary antibodies. Primary antibodies used were anti-V5 (mouse,

Thermo Scientific, Cat. # R960-25) and anti-GFAP (rabbit, Abcam, Cat. # ab7260). Secondary antibodies used were Alexa Fluor 546 goat anti-mouse secondary antibody (Jackson Immunoresearch, CAT# A-11003) and Cy3-AffiniPure Goat Anti-Rabbit secondary antibody (Jackson Immunoresearch, CAT# 111-165-008).

For brain histology, mice were anesthetized with an overdose of pentobarbital and performed intracardiac perfusion with phosphate buffer saline (PBS), followed by 4% PFA. Brains were removed and post-fixed in 4% PFA for 4 hr at RT. The samples were cryoprotection in 30% sucrose solution until they sank. For immunostaining, coronal sections were prepared on a Cryostat microtome (Leica CM1950), blocked with 3% BSA in PBS with 0.3% Triton X-100 for 1 hr, and followed by staining with the anti-GFAP (rabbit, Abcam, Cat. # ab7260) antibody at 4 ℃ overnight. After washing with PBS, the sections were incubated with 647 AffiniPure Goat Anti-Rabbit secondary antibody (Jackson Immunoresearch, CAT# 111-605-144) for 1 h at room temperature. PBS-washed sections were then coverslipped with 50% glycerol mounting medium. To observe AAV-mediated expression in the brain post-mortem, the sections (50 μm) were directly coverslipped with 50% glycerol mounting medium.

Fluorescent images were acquired on a Zeiss confocal microscope (LSM880, 40X, NA1.3 oil-immersion lens) or an Olympus VS120 virtual microscopy slide scanning system (10X objective) and further processed using Image J (NIH).

**Immunoblotting**

For western blotting, cOpn5-HEK 293T cells were pretreated with 10 μM staurosporine or buffer control for 60 min before irradiation with 470 nm light for 3 min (5s on, 5s off, 100 μW /mm$^2$). The protein samples were directly lysed into 1x SDS loading buffer immediately after the light was off. After electrophoresed by SDS–PAGE at 110 V, the separated proteins were transferred onto PVDF membranes and blocked with 5% nonfat milk in Tris-buffered with 0.1% Tween® 20 Detergent (TBST) buffer. Membranes were incubated with primary antibodies anti-Phospho-MARCKS (Cell Signaling Technology, CAT# 8722) and anti-alpha-tubulin (Sigma-Aldrich, CAT# T5168) in blocking buffer overnight. Membranes were washed three times in TBST before incubation in 1:30,000 species-specific HRP-conjugated antibodies (Thermo Fisher Scientific, Cat. # 8722 and Cat. # T5168). Membranes were washed again three times in TBST before visualization with Enhanced Chemiluminescence substrate. ImageJ was used to quantify protein bands from western blot films.

**cAMP and IP$_1$ Assay**

For IP$_1$ assays, HEK 293T cells that stably expressed cOpn5 or hOPN5 were optically stimulated for 3 min (100 μW /mm$^2$, 470 nm) min in LiCl-containing buffer, with or without 10nM YM-254890 pretreatment to block the G$_q$ proteins. After incubation (1 hour), accumulation of IP$_1$ was measured by a competitive immunoassay according to the manufacturer’s instructions with an ELISA kit (CisBio, Cat. # 72IP1PEA).

For cAMP assays, HEK 293T cells that stably expressed chicken, mouse, human Opn5s were incubated with or without 10 μM all trans-retinal 24 hr in the dark, and the cells were supplied with 10 μM forskolin 30 min before light stimulation with 470 nm for 3 min (5s on, 5s off, 100 μW /mm$^2$). The cAMP concentration in the cell lysis was measured with an ELISA kit according to the manufacturer's protocol (Wuhan Bioyeargene biotechnology, Cat. # BK101).

**$Ca^{2+}$ imaging of cell cultures**

Supplementary Table 2 provides detailed information on light excitation sources, microscopes, and key experimental parameters.

Specifically, HEK 293T cells were transiently expressed Opn5s (chicken, human and mouse) and loaded with $Ca^{2+}$ indicator Calbryte 630-AM (5μM) for 1 hr in HBSS buffer. Images were captured and analyzed by Nikon confocal microscopy (A1R, 25X, NA1.1 water-immersion lens) with the NIS-Elements software (Nikon). For image acquisition, we used the 488 nm from microscope light source.

cOpn5-HEK 293T stable cells were loaded with $Ca^{2+}$ indicator Calbryte 630-AM (5μM) for 1 hr in Hank's Balanced Salt Solution (HBSS). Nominal $Ca^{2+}$-free buffer was made by discarding the calcium in HBSS. Spinning disk confocal system was used to investigate the spectral characteristic of cOpn5. We adjusted the wavelength (365 nm, 395 nm, 470 nm, 515 nm, 561 nm, 590 nm and 630 nm), duration (1 ms, 5 ms, 10 ms, 20 ms and 50 ms; 470 nm), and intensity (0, 4.8, 8, 16, or 32 μW/mm$^2$; 470 nm; 10 ms) of optical stimulation.

We stimulated HEK 293T cells transiently expressing opto-a1AR with 515 nm light (7 mW/mm$^2$, 60 s). For hOPN4-expressing cells, we used 470 nm light (40 mW/mm$^2$, 25 s) with or without 10 μM All-trans-Retinal (ATR). For chemogenetic activation, we stimulated HEK 293T cells expressing hM3Dq with local puff applications of clozapine-N-oxide (CNO,100 nM) near the imaging field.

Primary astrocytes were infected with the *AAV2/8 GfaABC1D-cOpn5-T2A-eGFP.* After cOpn5 expression, astrocytes were loaded with the Calbryte™ 630-AM and stimulated with 488 nm light from microscope.

To observe the effect of spatially restricted stimulation on $Ca^{2+}$ signals, we used Nikon A1R enabled spatially patterned illumination and a 25×, NA1.1 water-immersion lens to stimulate a small area (488 nm; 3.7×3.7 μm$^2$ for stimulating single HEK 293T cells and 8.1×8.1 μm$^2$ for stimulating single astrocytes) to determine $Ca^{2+}$ wave propagations. The sampling rate was 2 fps at a size of 384×286 pixels. To observe $Ca^{2+}$ wave propagation within individual astrocytes, we used a Zeiss LSM 880 confocal microscope equipped spatially patterned illumination function and a 10×（NA 0.45) dry objective lens to stimulate a small area (4 × 4 μm$^2$) using blue light (488 nm) and imaged an area of 512×512 pixels at the frame rate of 3.3 fps.

Acquired data were analyzed by the Matlab and ImageJ software. Typically, we photostimulated and imaged 5 separate regions per well and repeated 3 wells per condition.

**Behavioral tasks and *in vivo* optogenetics**

Mice had *ad libitum* access to normal mouse lab pellet food and water before the tests. Mice were allowed to recover for at least two weeks following stereotaxic infusion of AAV vectors and the implant of optic fiber ferrule. On the test day, an optical fiber (200 μm diameter, 0.37 NA; Shanghai Fiblaser) was connected to the ferrule. *VGAT*-Cre mice expressing cOpn5 and EGFP or EGFP alone were placed in the test chamber. To generate light pulses, we used a Master-8 pulse stimulator (A.M.P.I., Israel) to control the light delivery (473 nm; peak light power 0.75 mW) from a solid state laser (MBL-III-473, Changchun New Industries Optoelectronics Technology, China). For the food intake assay, we measured the amount of food consumed before and during optogenetic stimulation (30s on, 30 s off using 20 Hz blue light pulses) that lasted for one hour. For the food foraging test, high fat food pellets were hidden in the bedding litter. Mouse received light stimulation until the hidden food pellet was retrieved. Foraging time was calculated since the blue light on until the mouse got the food. Mouse behaviors were videotaped via a camera 30 cm above the chamber. Pellet retrieval time was calculated by manual video scoring offline.

**Electrophysiological recording and $Ca^{2+}$ Imaging in brain slices**

Slice preparation and electrophysiological recordings were performed as described previously [2]. Briefly, mice were deeply anesthetized with pentobarbital (100 mg/kg i.p.) and intracardially perfused with 5 mL ice-cold oxygenated perfusion solution containing (in mM) 1.25 glucose, 225 sucrose, 119 NaCl, 2.5 KCl, 0.1 $CaCl_2$, 4.9 $MgCl_2$, 1.0 $NaH_2PO_4$, 26.2 $NaHCO_3$, 3 kynurenic acid, and 1 Na L-ascorbate (all chemicals for brain slicing recording were from Sigma-Aldrich USA). The mouse brain was rapidly removed and placed in ice-cold oxygenated slicing solution containing (in mM) 110 choline chloride, 2.5 KCL, 0.5 $CaCl_2$, 7 $MgCl_2$, 1.3 $NaH_2PO_4$, 25 $NaHCO_3$, 20 Glucose, 1.3 Na ascorbate, and 0.6 Na pyruvate. Coronal sections (250 μm thick) were prepared with a Leica VT1200S vibratome. The slices were incubated for 1 h at 34°C with ringer solution saturated with a 95%:5% mixture of $O_2$: $CO_2$. The ringer solution consisted of (in mM) 125 NaCl, 2.5 KCl, 2 $CaCl_2$, 1.3 $MgCl_2$, 1.3 $NaH_2PO_4$, 25 $NaHCO_3$, 10 Glucose, 1.3 Na ascorbate, and 0.6 Na pyruvate. The intrapipette solution contained (in mM) 130 K-gluconate, 10 HEPES, 0.6 EGTA, 5 KCl, 3 $Na_2ATP$, 0.3 $Na_3GTP$, 4 $MgCl_2$, and 10 $Na_2$-phosphocreatine (pH 7.2–7.4). Whole-cell patch-clamp recordings were made on mouse brain slices using a Multiclamp 700B amplifier, which was connected to a Digidata 1440 digitizer (Molecular Devices) and attached to a PC running pClamp 10. Electrophysiological data were acquired at 5 kHz and low-pass filtered at 10 kHz.

For $Ca^{2+}$ imaging, the red fluorescent signals from Calbryte™ 630 were imaged on a two-photon microscope (Olympus BX61W1, 20×, NA1.1 water-immersion lens, 2 fps). For photostimulation, blue light pulses (470 nm) were generated with an LED.

**Two-photon imaging of neuronal $Ca^{2+}$ signals after astrocytes activation *in vivo***

For *in vivo* two-photon imaging of neuronal $Ca^{2+}$ signals, 600 nl AAV vectors separately expressing jGCaMP7b and cOpsin5 were mixed and injected into the S1 cortex of adult (>6 weeks of age) C57BL/6N mice. Open skull surgery and microscopy device were the same as the imaging of ATP release. Two photon imaging was performed ~200 μm below the dura mater (layer 2/3). The laser excitation wavelength of jGCaMP7b was 920 nm. For neuronal calcium imaging, the scanning was set at 5 fps, ×1 optical zoom and 512×512-pixel resolution.

To process and analyze imaging data, MATLAB R2018a and Fiji (NIH) were used. Lateral shifts were corrected using the Image Stabilizer plugin of Fiji. The $Ca^{2+}$ signals and decoded spike rates were analyzed using MATLAB program package, CNMF_E-master. Relative spike rate was normalized to average basal of all neurons (n = 193). The statistics of $Ca^{2+}$ events per min and spike rate were performed using Prism 9.

**Two-photon imaging of ATP release *in vivo***

For *in vivo* two-photon imaging of ATP release, we mixed the AAV vectors for expressing the $GRAB_{ATP}$ sensor and GfaABC1D-cOpn5-T2A-mCherry, and then infused 600 nl of the AAV mixtures into the layer 1 of S1 cortex of adult (>6 weeks of age) C57BL/6N mice using a nanoliter injector. Two weeks after vector infusion, a circular skull area of 3 mm in diameter was drilled over the virus injection site and covered by a glass window. A stainless steel head-fixation bar that included an imaging chamber was positioned over the window and affixed to the exposed skull with dental cement. Mice were allowed to recover from the surgery for a week. Two-photon imaging was performed ~100 µm below the dura mater using a FluoView FVMPE-RS microscope (Olympus, 25×, NA1.05 water-immersion lens) equipped with a pulsed laser (Spectra-Physics). The laser wavelength for activating cOpn5 and exciting the fluorescent $GRAB_{ATP}$ imaging was 920 nm, and that for validating the mCherry-expression was 1045 nm. For imaging $GRAB_{ATP}$ signals, we used 1.5 fps, ×1 optical zoom, and 512×512-pixel sampling. The same setting was used in no-cOpn5 and cOpn5 groups. Lateral shifts across image frames were corrected using the TurboReg plugin of Fiji (NIH). ATP release events were identified and quantified using the Astrocyte Quantitative Analysis (AQuA) software [3], which took an event-based perspective to model and quantify ATP release in the form of flashes. Every single burst event was pseudo-colored in the imaging field and the heat maps of *ΔF/F* were plotted using MATLAB R2021a and analyzed using Prism9. The statistics of ATP release numbers were taken using Prism 9. For CNO-treated mice (hM3Dq expression), mice received a single dose of CNO (2 mg/kg) by intraperitoneal injection 40 min prior to imaging.

**$Ca^{2+}$ signal analysis**

The imaging images were converted into TIF files and analyzed by CaImAnalyzer MATLAB program. F0 is the average value of the first 5 seconds as the basal fluorescence value. DeltaF is the fluorescence value of each frame minus F0. DeltaF /F0 values are stored in CalData MATLAB files，then use group_data_plot MATLAB program plot curve.

**$Ca^{2+}$ signal propagation analysis**

The imaging images were converted into TIF files and analyzed by CellWaveforCell MATLAB program. The deltaF/F0 image results were obtained after running. F0 is the average value of the first 10 frames as the basal fluorescence value. DeltaF is the fluorescence value of each frame minus F0. The results of image deltaF/F0 were opened with Fiji, and the pseudo-color image of fluorescence signal changing with time was obtained by image, Hyperstacks and grey-color code.

### Statistics

We used MATLAB 2018a and GraphPad Prism 9 to perform statistical analysis. Statistical method, the corresponding p values, and the sample sizes are reported in the figure legends as well as in Supplementary Table 3. The sample sizes were not predetermined. In all optogenetic activation experiments and fiber photometry recordings, sample sizes (n) denote the number of stimulation or recording sites. In slice recording experiments, sample sizes denote the cell numbers recorded. Data were reported as mean ± SEM in all Figures.

### Data and Code availability

Data and custom programs are available upon request. All the programs used are on the institute hosted server and can be accessed at http://imaging.cibr.ac.cn/Public/Upload/Content/20211025/617639f0637c2.rar.


### ACKNOWLEDGMENTS

This work was supported by Beijing Municipal Government.


### Author information

These authors contributed equally: Ruicheng Dai, Tao Yu, Danwei Weng

**CONTRIBUTIONS:**

R.C.D, T.Y, D.W.W, and M.L. designed the research and wrote the manuscript. H.L performed Two-photon imaging of ATP release. Y.T.C and Y.Q.R performed electrophysiological recording. Z.F.W. and Y.L.L provided ATP sensor. Q.C.G, X.W.G, and Z.Y.Q analyzed $Ca^{2+}$ signal. H.Y.Z and W.T.W performed behavior experiments. W.T.W performed sequence analysis. Q.C.G and S.W. set up the LED stimulation system.

### Supplementary ReferencesUncategorized References


1 Schildge, S., Bohrer, C., Beck, K. & Schachtrup, C. Isolation and culture of mouse cortical astrocytes. *Journal of visualized experiments: JoVE* (2013).

2 Zhang, J. *et al.* Presynaptic excitation via GABAB receptors in habenula cholinergic neurons regulates fear memory expression. *Cell* **166**, 716-728 (2016).

3 Wang, Y. *et al.* Accurate quantification of astrocyte and neurotransmitter fluorescence dynamics for single-cell and population-level physiology. *Nature neuroscience* **22**, 1936-1944 (2019).
