## Supplementary Table 1-4 for "A Neuropsin-based Optogenetic Tool for Precise Control of G_q_ signaling"

### Supplementary Table 1: Comparison of chicken, human, and mouse Opn5

| Opsins | Alias | Species | Sources |
|---|---|---|---|
| Chicken Opn5 | cOpn5 | Chelonia mydas | GenBank NM_001130743.1 |
| Human OPN5 | hOPN5 | Homo sapiens | GenBank AY377391.1 |
| Mouse Opn5 | mOpn5 | Mus musculus | GenBank NM_181753.4 |
| Mouse Opn4L | mOpn4L | Mus musculus | GenBank NM_013887.2 |

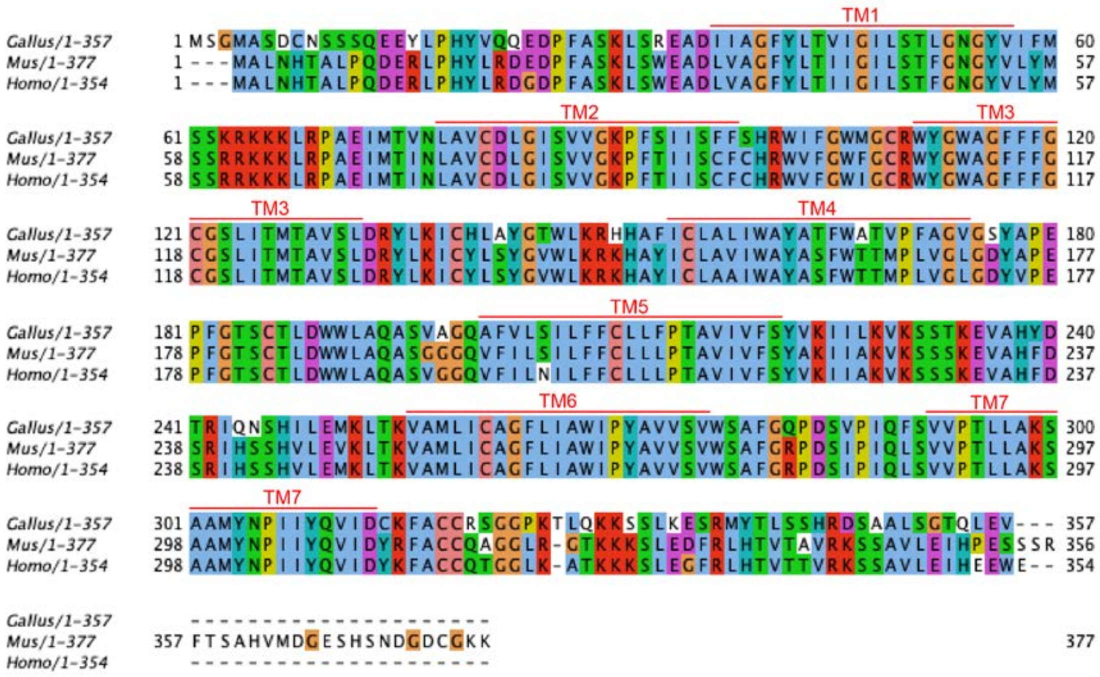

**Supplementary Table 2: Key resources**

| | |
|---|---|
| V5-cOpn5 forward primer | 5’-cgtgaggtaccggatcctctagaatgggcaagcccatccccaaccccctgctgg gcctggacagcaccatgagtgggatggcatcggac-3’ |
| V5-cOpn5 reverse primer | 5’-tcgataagcttgatatcgaattcttagacttccagttgggttccgct-3’ |
| cOpn5-T2A-eGFP hSyn promoter forward primer | 5’-tagagtcgagctcaagcttgccaccatgagtgggatggcatcggactgca-3’ |
| cOpn5-T2A-eGFP hSyn promoter reverse primer | 5’-aaccgcgggccctctagagcatatgttacttgtacagctcgtccatgccg-3’ |
| cOpn5-T2A-eGFP GfaABC1D promoter forward primer | 5’-acctccgctgctcgcggggtctagaatgagtgggatggcatcggactgca-3’ |
| cOpn5-T2A-eGFP GfaABC1D promoter reverse primer | 5’-tatcgataagcttgatatcgaattcttacttgtacagctcgtccatgccg-3’ |
| cOpn5-T2A-eGFP EF1a promoter forward primer | 5’-tacattatacgaagttatggcgcgccttattacttgtacagctcgtccatg-3’ |
| cOpn5-T2A-eGFP EF1a promoter reverse primer | 5’-atactttatacgaagttatgctagccaccatgagtgggatggcatcggactg-3’ |
| cOpn5-T2A-mCherry forward primer | 5’-gcatcacctccgctgctcgcggggtatgagtgggatggcatcggactgca -3’ |
| cOpn5-T2A-mCherry reverse primer | 5’-tcaccatggtggcgaccgggggatctgggccaggattctcctcgacgtca -3’ |
| pcDNA3.1-opto-a1AR-EYFP | Addgene plasmid #20947 |
| EGFP-CAAX | Gift from Yulong Li |
| pLJM1-EGFP | Addgene plasmid #19319 |
| pAAV-GfaABC1D-hM3D(Gq)-mCherry | Addgene Plasmid #50478 |
| pAAV-EF1a-DIO-eGFP-WPRE-pA | This study |
| PAAV-hSyn-GOI | This study |
| pLJM1-cmv-cOpn5 | This study |
| pLJM1-cmv-hOPN5 | This study |
| pLJM1-cmv-mOpn5 | This study |
| pLJM1-cmv-V5-Opn5 | This study |
| pLJM1-cmv-cOpn5-T2A-eGFP | This study |
| PAAV-hSyn-cOpn5-T2A-eGFP-WPR-pA | This study |
| PAAV-GfaABC1D-cOpn5-T2A-eGFP-WPR-pA | This study |
| pAAV-EF1a-DIO-cOpn5-T2A-eGFP-WPRE-pA | This study |
| PAAV-GfaABC1D-cOpn5-T2A-mCherry-WPR-pA | This study |
| Lenti-cmv-cOpn5-puro | Chinese Institute for Brain Research, Beijing |
| Lenti-cmv-hOPN5-puro | Chinese Institute for Brain Research, Beijing |

| | |
|---|---|
| Lenti-cmv-mOpn5-puro | Chinese Institute for Brain Research, Beijing |
| Lenti-cmv- hM3Dq -puro | Chinese Institute for Brain Research, Beijing |
| AAV2/9-EF1a-DIO-cOpn5-T2A-eGFP | Chinese Institute for Brain Research, Beijing |
| AAV2/9-hSyn-cOpn5-T2A-eGFP | Chinese Institute for Brain Research, Beijing |
| AAV2/9-Ef1a-DIO-cOpn5-T2A-eGFP | Chinese Institute for Brain Research, Beijing |
| AAV2/8-GFaABC1D-cOpn5-T2A-eGFP | Chinese Institute for Brain Research, Beijing |
| AAV2/8-GfaABC1D-cOpn5-T2A-mCherry | Chinese Institute for Brain Research, Beijing |
| AAV2/9-EF1a-EGFP | Chinese Institute for Brain Research, Beijing |
| AAV2-EF1α-DIO-GCaMP6m | Chinese Institute for Brain Research, Beijing |
| AAV2/9-GFaABC1D-GCaMP6s | Chinese Institute for Brain Research, Beijing |
| AAV2/9-GfaABC1D-ATP1.0 | WZ Biosciences Inc. Cat. # YL006003-AV9 |
| AAV9-hSyn-NES-jRGECO1a-WPRE | WZ Biosciences Inc. Cat. # BS8-NOAAAV9 |
| AAV2/9-mCaMKIIa-jGCaMP7b-WPRE-pA | Shanghai Taitool Bioscience Co., Ltd Cat. # S0712-9-H20 |
| Light excitation sources in Figs 1c, 1d, 1e, 1f, 2d, 4e | Nikon A1R MP 488 nm |
| Light excitation sources in Figs 1g, 2a, 2b, 2c, 2h, 2i, 4a, 4b and Extended Figs 1a,1b, 1c, 1d, 1e, 2a | Thorlabs M470L3 470 nm mounted LED |
| Light excitation sources in Figs 3b, 3c | Olympus BX61W1 Two-photon microscope 488 nm |
| Light excitation sources in Fig 4d | Zeiss LSM 880 Confocal laser scanning microscope 488 nm |
| Light excitation sources in Fig 2a | LG3535 365 nm mounted LED wavelength coverage: 360-370 nm |
| Light excitation sources in Fig 2a | LG3535 395 nm mounted LED wavelength coverage: 390-400 nm |
| Light excitation sources in Fig 2a | 561 nm laser from Changchun New Industries Optoelectronics Technology, China MGL-FN-561 |
| Light excitation sources in Fig 2a | 590 nm mounted LED wavelength coverage: 570-615 nm |
| Light excitation sources in Fig 2a | 630 nm mounted LED wavelength coverage: 615-660 nm |
| Light excitation sources in Fig 2f, 2g | 515 nm laser Changchun New Industries Optoelectronics Technology, China, MGL-F-515 |
| Microscope for Figs 1c, 1d, 1e, 1f, 2d, 4e and Extended Fig 1a | Nikon A1R MP Multiphoton confocal microscopes |

| | |
|---|---|
| Microscope for Figs 2a, 2b, 2c, 2f, 2g, 2h, 2i, 4a, 4b and Extended Figs 1a | Nikon ECLIPASE Ti Spinning Disk |
| Microscope for Figs 3b, 3c and Extended Figs 2a, 2b, 2c | Olympus BX61W1 Two-photon microscope |
| Microscope for Figs 1b, 4d | Zeiss LSM 880 Confocal laser scanning microscope |
| Microscope for Fig 4g, 4h, 4i, 4j, 5 and Extended Fig 4 | FluoView FVMPE-RS Multiphoton microscope |

### Supplementary Table 3: Summary of statistical analyses

| Fig | Conditions | n per group | Analysis | P value |
|---|---|---|---|---|
| 1f | cOpn5 group:<br>light vs YM-254890 | 19, 15 | one way ANOVA | P < 0.0001 |
| 1g | ctrl vs. light | 7,6 | Unpaired *t* test | P = 0.0005 |
| | light vs.<br>light+ YM-254890 | 6,5 | Unpaired *t* test | P = 0.0004 |
| 3e | $LH^{cOpn5}$ group:<br>light on vs. off | 5, 5 | Unpaired *t* test | P =0.0003 |
| | $LH^{EGFP}$ group:<br>light on vs. off | 5, 5 | Unpaired *t* test | P =0.3466 |
| 3g | $ZI^{cOpn5}$ group:<br>light on vs. off | 5, 5 | Unpaired *t* test | P < 0.0001 |
| | $ZI^{EGFP}$ group:<br>light on vs. off | 4, 4 | Unpaired *t* test | P =0.7608 |
| 3i | LH group:<br>ctrl vs. light | 6, 6 | Unpaired *t* test | P =0.00041 |
| | ZI group:<br>ctrl vs. light | 5, 5 | Unpaired *t* test | P =0.0027 |
| 5g | ctrl vs. hM3Dq+CNO | 6,5 | Unpaired *t* test | P =0.5790 |
| | ctrl vs. cOpn5 | 6,5 | Unpaired *t* test | P =0.0003 |
| | hM3Dq+CNO vs. cOpn5 | 5,5 | Unpaired *t* test | P =0.0009 |
| 5j | cOpn5 spike rate:<br>0-5 min vs. 15-20 min | 193, 193 | Unpaired *t* test | P < 0.0001 |
| | cOpn5 $Ca^{2+}$ frequency:<br>0-5 min vs. 15-20 min | 193, 193 | Unpaired *t* test | P < 0.0001 |
| 5k | mCherry Spike rate:<br>0-5 min vs. 15-20 min | 170, 170 | paired *t* test | P =0.0164 |
| | mCherry $Ca^{2+}$ frequency:<br>0-5 min vs. 15-20 min | 170, 170 | paired t test | P =0.4659 |
| Extended 1c | ctrl vs. light | 4, 4 | Tukey's multiple comparisons test | P =0.0096 |
| | light vs.<br>light+ staurosporine | 4, 4 | Tukey's multiple comparisons test | P = 0.0004 |
| Extended 1d | ctrl vs. light | 4,4 | Unpaired *t* test | P =0.4338 |
| Extended 1e-left | ctrl vs. light | 3,3 | Unpaired *t* test | P =0.992 |
| Extended 1e-Right | cOpn5 group:<br>ctrl vs. light | 4,4 | Sidak's multiple comparisons test | P =0.0223 |
| | hOPN5 group:<br>ctrl vs. light | 4,4 | Sidak's multiple comparisons test | P< 0.0001 |
| | mOpn5 group:<br>ctrl vs. light | 4,4 | Sidak's multiple comparisons test | P< 0.0001 |
| Extended 4c | Basal vs. light | 29, 29 | Unpaired *t* test | P < 0.0001 |

### Supplementary Table 4: Comparison cOpn5 with other optogenetic tools

| Tools and references | Wavelength $\lambda_{max}$ (nm) | Light sensitivity | Stimulation duration | exogenous chromo-phores | Response amplitude | Tested cells |
|---|---|---|---|---|---|---|

| | | | | | | |
|---|---|---|---|---|---|---|
| cOpn5<br>this study | 470 nm | 16 μW/$mm^2$ | 10 ms | No | Peak Ca2+ responses 3.0-8.0 (*ΔF/F*) | HEK 293T cells astrocytes, and neurons |
| opto-α1AR [1] | 500 nm | 7 mW/$mm^2$ | 60 s | No | peak $Ca^{2+}$ response ~0.23 (*ΔF/F*) | HEK 293 cells |
| opto-α1AR [2] | 473 nm | 20 Hz, 45-ms light pulses, 5 mW | 5 min | No | >20 % increase in sIPSC frequency | astrocytes |
| opto-α1AR [2] | 510 nm | 7 mW/$mm^2$ | 60 s | No | peak $Ca^{2+}$ response ~0.5 (*ΔF/F*) | HEK 293T cells |
| mouse melanopsin (Opn4) [3] | 480 nm | $10^{15}$ photons $s^{-1}$ $cm^{-2}$ at 500 nm; ~40 mW/$mm^2$ | >60 s | 11-cis-retinal dehyde | peak $Ca^{2+}$ response ~0.1 (ΔF/F) | HEK293-TRPC 3 cells |
| mouse melanopsin and its mutants [4] | 488 nm | a white fluorescent light source (intensity undefined) | 60 s | 11-cis retinal | peak $Ca^{2+}$ response ~0.25 (ΔF/F) for the best mutant $Opn4^{9A}$ | CHO cells |
| Human melanopsin (OPN4) [5] | 473 nm | 7 mW/$mm^2$ | 20 s | Unknown | no significant change in $Ca^{2+}$ amplitude; about 1 fold increase in $Ca^{2+}$ event frequency | *in vivo* astrocytes |
| human melanopsin [5] | 470 nm | 40 mW/$mm^2$ | 25 s | ATR | peak $Ca^{2+}$ response ~0.65 (ΔF/F) | HEK 293T cells |
| Wild-type ChR2 [6,7] | 470 nm | 8–12 mW/$mm^2$ | 2.3 ± 1.1 ms | No | 731± 100 pA with steady state/peak current ratio: 0.4 ± 0.04 | hippocampal cell culture |
| ChR2 H134R [8] | 450 nm | ~10 mW/$mm^2$ (470 nm) | 0.96 ± 0.12 ms | No | 4.47 nA | HEK 293T cells |
| ChETA [9] | 490 nm | ~10 mW/$mm^2$ | 0.9 ± 0.1 ms | No | 645 ± 47 pA with steady state/peak current ratio: 0.6 ± 0.04 | hippocampal cell culture |
| ChrimsonR [10] | 590 nm | 4.6 mW/$mm^2$ | 0.9 ± 0.1 ms | No | ~300 pA | cultured neurons |
